# The discovery, distribution and diversity of DNA viruses associated with *Drosophila melanogaster* in Europe

**DOI:** 10.1101/2020.10.16.342956

**Authors:** Megan A. Wallace, Kelsey A. Coffman, Clément Gilbert, Sanjana Ravindran, Gregory F. Albery, Jessica Abbott, Eliza Argyridou, Paola Bellosta, Andrea J. Betancourt, Hervé Colinet, Katarina Eric, Amanda Glaser-Schmitt, Sonja Grath, Mihailo Jelic, Maaria Kankare, Iryna Kozeretska, Volker Loeschcke, Catherine Montchamp-Moreau, Lino Ometto, Banu Sebnem Onder, Dorcas J. Orengo, John Parsch, Marta Pascual, Aleksandra Patenkovic, Eva Puerma, Michael G. Ritchie, Omar Rota-Stabelli, Mads Fristrup Schou, Svitlana V. Serga, Marina Stamenkovic-Radak, Marija Tanaskovic, Marija Savic Veselinovic, Jorge Vieira, Cristina P. Vieira, Martin Kapun, Thomas Flatt, Josefa González, Fabian Staubach, Darren J. Obbard

## Abstract

*Drosophila melanogaster* is an important model for antiviral immunity in arthropods, but very few DNA viruses have been described from the family Drosophilidae. This deficiency limits our opportunity to use natural host-pathogen combinations in experimental studies, and may bias our understanding of the *Drosophila* virome. Here we report fourteen DNA viruses detected in a metagenomic analysis of approximately 6500 pool-sequenced *Drosophila*, sampled from 47 European locations between 2014 and 2016. These include three new Nudiviruses, a new and divergent Entomopox virus, a virus related to *Leptopilina boulardi* filamentous virus, and a virus related to *Musca domestica* salivary gland hypertrophy virus. We also find an endogenous genomic copy of Galbut virus, a dsRNA Partitivirus, segregating at very low frequency. Remarkably, we find that *Drosophila* Vesanto virus, a small DNA virus previously described as a Bidnavirus, may be composed of up to 12 segments and represent a new lineage of segmented DNA viruses. Two of the DNA viruses, *Drosophila* Kallithea nudivirus and *Drosophila* Vesanto virus are relatively common, found in 2% or more of wild flies. The others are rare, with many likely to be represented by a single infected fly. We find that virus prevalence in Europe reflects the prevalence seen in publicly-available datasets, with *Drosophila* Kallithea nudivirus and *Drosophila* Vesanto virus the only ones commonly detectable in public data from wild-caught flies and large population cages, and the other viruses being rare or absent. These analyses suggest that DNA viruses are at lower prevalence than RNA viruses in *D. melanogaster*, and may be less likely to persist in laboratory cultures. Our findings go some way to redressing an earlier bias toward RNA virus studies in *Drosophila*, and lay the foundation needed to harness the power of *Drosophila* as a model system for the study of DNA viruses.

## Introduction

*Drosophila melanogaster* is one of our foremost models for antiviral immunity in arthropods (Huszart and Imler 2008, Mussabekova *et al*. 2017) and more than 100 *Drosophila-associ*ated viruses have been reported, including at least 30 that infect *D. melanogaster* (Brun and Plus 1980, Wu *et al*. 2010, Longdon *et al*. 2015, Webster *et al*. 2015, Webster *et al*. 2016, Medd *et al*. 2018). These include viruses with positive sense single-stranded RNA genomes (+ssRNA), such as *Drosophila* C virus, negative sense RNA genomes (-ssRNA), such as *Drosophila* melanogaster sigmavirus, and double-stranded RNA genomes (dsRNA), such as Galbut virus. Many of these viruses are common in laboratory fly cultures and in the wild (Webster *et al*. 2015). For example, the segmented and vertically-transmitted Galbut virus is carried by more than 50% of wild-caught adult *D. melanogaster* (Webster *et al*. 2015, Cross *et al*. 2020). Overall, more than 20% of wild-caught flies carry multiple RNA viruses, and about one third of laboratory fly lines and almost all *Drosophila* cell cultures are infected by at least one RNA virus (Plus 1978, Brun and Plus 1980, Webster *et al*. 2015, Shi, White, *et al*. 2018). However, in contrast to this wealth of RNA viruses, DNA viruses of *Drosophila* were unknown until relatively recently (Brun and Plus 1980, Huszart and Imler 2008).

The first described DNA virus of a drosophilid was published only ten years ago, after discovery through metagenomic sequencing of wild-caught *Drosophila innubila* (Unckless 2011). This virus is a member the Nudiviridae, a lineage of large (120-180Kbp) dsDNA viruses that are best known as pathogens of Lepidoptera and Coleoptera (Harrison *et al*. 2020), but which have genomic ‘fossil’ evidence across a broad host range (Cheng *et al*. 2020). *Drosophila innubila* Nudivirus infects several *Drosophila* species in North America, with a prevalence of up to 40% in *D. innubila*, where it can substantially reduce fecundity and lifespan (Unckless 2011, Hill and Unckless 2020). The first reported DNA virus of *D. melanogaster* was a closely-related Nudivirus reported by Webster *et al*. (2015), and provisionally called ‘Kallithea virus’ after a collection location. This virus was also first detected by metagenomic sequencing, but PCR surveys indicate that it is common in wild *D. melanogaster* and *D. simulans* (globally 5% and 0.5% respectively; Webster *et al*. 2015). *Drosophila* Kallithea nudivirus has been isolated for experimental study, and reduces male longevity and female fecundity (Palmer *et al*. 2018). Consistent with its presumed niche as a natural pathogen of *Drosophila*, this virus encodes a suppressor of *D. melanogaster* NF-kappa B immune signalling (Palmer *et al*. 2019). Prior to the work described here, the only other reported natural DNA virus infection of a drosophilid was the discovery— again through metagenomic sequencing—of a small number of RNA reads from Invertebrate iridescent virus 31 (IIV31; *Armadillidium vulgare* iridescent virus) in *D. immigrans* and *D. obscura* (Webster *et al*. 2016). This virus is known as a generalist pathogen of terrestrial isopods (Piegu *et al*. 2014), but its presence as RNA (indicative of expression) in these *Drosophila* species suggests that it may have a broader host range.

The apparent dearth of specialist DNA viruses infecting *Drosophilidae* is notable (Brun and Plus 1980, Huszart and Imler 2008), perhaps because DNA viruses have historically dominated studies of insects such as Lepidoptera (Cory and Myers 2003), and because DNA viruses are well known from other Diptera, including the Hytrosaviruses of *Musca* and *Glossina* (Kariithi *et al*. 2017), Densoviruses of mosquitoes (Carlson *et al*. 2006), and Entomopox viruses of various Culicomorpha (Lawrence 2011). The lack of native DNA viruses for *D. melanogaster* has practical implications for research, as the majority of experiments have had to utilise viruses that do not naturally infect *Drosophila*, and which have not co-evolved with them (Bronkhorst *et al*. 2014, West and Silverman 2018, but see Palmer *et al*. 2019).

It not only remains an open question as to whether the *D. melanogaster* virome is really depauperate in DNA viruses; the prevalence of DNA viruses in *Drosophila*, their phylogenetic diversity, their spatial distribution and temporal dynamics, and their genetic diversity all remain almost unstudied. However, many of these questions can be addressed through large-scale metagenomic sequencing of wild-collected flies. As part of a large *Drosophila* population genomics study using pool-sequencing of wild *D. melanogaster*, we previously reported the genomes of four DNA viruses associated with European *Drosophila* samples collected in 2014 (the DrosEU consortium; Kapun *et al*. 2020). These included a second *melanogaster*-associated Nudivirus (there referred to as ‘Esparto virus’), two Densoviruses (‘Viltain virus’ and ‘Linvill road virus’), and two segments of a putative Bidnavirus (‘Vesanto virus’). Here we expand our sampling to encompass 167 shortread pool-sequenced samples from a total of 6668 flies, collected seasonally over three years from 47 different locations across Europe. We use these population genomic data as a metagenomic source to discover additional DNA viruses, estimate their prevalence in time and space, and quantify levels of genetic diversity.

We complete the genome of a novel and highly divergent Entomopox virus, identify a further three *Drosophila-associated* Nudiviruses, fragments of a novel Hytrosavirus most closely related to *Musca domestica* salivary gland hypertrophy virus, fragments of a Filamentous virus distantly related to *Leptopilina boulardi* filamentous virus, and three polinton-like sequences related to ‘Adintoviruses’. Our improved assemblies and sampling show that Vesanto virus may be composed of up to 12 segments, and appears to represent a new distinct lineage of multi-segmented ssDNA viruses related to the Bidnaviridae. We find that two viruses (*Drosophila* Kallithea nudivirus and *Drosophila* Vesanto virus) are common in European *D. melanogaster*, but that the majority of DNA viruses appear very rare—most probably appearing once in our sampling.

## Methods

### Sample collection and sequencing

A total of 6668 adult male *Drosophila* were collected across Europe by members of the DrosEU consortium between 19^th^ June 2014 and 22^nd^ November 2016, using yeast-baited fruit (Kapun *et al*. 2020, Kapun *et al*. 2021). There were a total of 47 different collection sites spread from Recarei in Portugal (8.4° West) to Alexandrov in Russia (38.7° East), and from Nicosia in Cyprus (36.1° North) to Vesanto in Finland (62.6° North). The majority of sites were represented by more than one collection, with many sites appearing in all three years, and several being represented by two collections per year (early and late in the *Drosophila* flying season for that location). After morphological examination to infer species identity, a minimum of 33 and maximum of 40 male flies (mean 39.8) were combined for each site and preserved in ethanol at −20°C or −80°C for pooled DNA sequencing. Male flies were chosen because, within Europe, male *D. melanogaster* should be morphologically unambiguous. Nevertheless, subsequent analyses identified the occasional presence of the sibling species *D. simulans*, and two collections were contaminated with the distant relatives *D. phalerata* and *D. testacea* (below). Full collection details are provided via figshare repository 10.6084/m9.figshare.14161250, and the detailed collection protocol is provided as supporting material in Kapun *et al* (2020).

To extract DNA, ethanol-stored flies were rehydrated in water and transferred to 1.5 ml well plates for homogenisation using a bead beater (Qiagen Tissue Lyzer II). Protein was digested using Proteinase K, and RNA depleted using RNAse A. The DNA was precipitated using phenol-chloroform-isoamyl alcohol and washed before being air dried and re-suspended in TE. For further details, see the supporting material in Kapun *et al* (2020). DNA was sequenced in three blocks (2014, most of 2015, 2016 and remainder of 2015) by commercial providers using 151nt paired end Illumina reads. Block 1 libraries were prepared using NEBNext Ultra DNA Lib Prep-24 and NEBNext Multiplex Oligos, and sequenced on the Illumina NextSeq 500 platform by the Genomics Core Facility of the University Pompeu Fabra (UPF; Barcelona, Spain). Block II and III libraries were prepared using the NEBNext Ultra II kit and sequenced on the HiSeq X platform by NGX bio (San Francisco, USA). All raw Illumina read data are publicly available under SRA project accession PRJNA388788.

To improve virus genomes, and following an initial exploration of the Illumina data, we pooled the remaining DNA from four of the collections (samples UA_Yal_14_16, ES_Gim_15_30, UA_Ode_16_47 and UA_Kan_16_57) for long-read sequencing using the Oxford Nanopore Technology ‘Minion’ platform. After concentrating the sample using a SpeedVac (ThermoFisher), we prepared a single library using the Rapid Sequencing Kit (SQK-RAD004) and sequenced it on an R9.4.1 flow cell, subsequently calling bases with Guppy version 3.1.5 (https://community.nanoporetech.com).

### Read mapping and identification of contaminating taxa

We trimmed Illumina sequence reads using Trim Galore version 0.4.3 (Krueger 2015) and Cutadapt version 1.14 (Martin 2011). To remove *Drosophila* reads, and to quantify potentially contaminating taxa such as *Wolbachia* and other bacteria, fungi, and trypanosomatids, we mapped each dataset against a combined *‘Drosophila* microbiome’ reference. This reference comprised the genomes of *D. melanogaster* (Chang and Larracuente 2019), *D. simulans* (Nouhaud 2018), three *Drosophila-asso*ciated *Wolbachia* genomes, 69 other bacteria commonly reported to associate with *Drosophila* (including multiple *Acetobacter*, *Gluconobacter*, *Lactobacillus*, *Pantoea*, *Providencia*, *Pseudomonas* and *Serratia* genomes), and 16 microbial eukaryotic genomes (including two *Drosophila*-associated trypanosomatids, a microsporidian, the entomopathogenic fungi *Metarhizium anisopliae*, *Beauveria bassiana* and *Entomophthora muscae*, and several yeasts associated with rotting fruit; the list of sequence accessions is provided in figshare repository 10.6084/m9.figshare.14161250). All mapping was performed using Bowtie 2 version 2.3.4 or version 2.4.1 (Langmead and Salzberg 2012) and we recorded only the best mapping position for each read, based on alignment match score (the Bowtie 2 default). To provide approximate quantification we used raw mapped read counts, normalised by target length and fly read counts where appropriate.

During manual examination of *de novo* assemblies (below) we identified a number of short contigs from other taxa, including additional species of *Drosophila, Drosophila* commensals such as mites and nematodes, and potential sequencing contaminants such as humans and model organisms. To quantify this potential contamination, we re-mapped all trimmed read pairs to a reference panel of short diagnostic sequences. This panel comprised a region of *Cytochrome Oxidase I* (COI) from 20 species of *Drosophila* (European *Drosophila* morphologically similar to *D. melanogaster*, and *Drosophila* species identified in *de novo* assemblies), 667 species of nematode (including lineages most likely to be associated with *Drosophila*, and a contig identified by *de novo* assembly), 106 parasitic wasps (including many lineages commonly associated with *Drosophila*), two species of mite (identified in *de novo* assemblies), complete mitochondrial genomes from six model vertebrates, and complete plastid genomes from eight crop species. Because crossmapping between *D. melanogaster* and *D. simulans* is possible at many loci, we also included a highly divergent but low-diversity 2.3 kbp region of the single-copy gene *Argonaute-2* to estimate levels of *D. simulans* contamination. Where reads indicated the presence of other *Drosophila* species, this was further confirmed by additional mapping to *Adh*, *Amyrel*, *Gpdh* and *6-PGD*. A full list of the reference sequences is provided in via figshare repository 10.6084/m9.figshare.14161250.

### Virus genome assembly and annotation

To identify samples containing potentially novel viruses, we retained read pairs that were not concordantly mapped to the combined ‘*Drosophila* microbiome’ reference (above) and used these for *de novo* assembly using SPAdes version 3.14.0 with the default spread of k-mer lengths (Nurk *et al*. 2013), after *in silico* normalisation of read depth to a target coverage of 200 and a minimum coverage of 3 using bbnorm (https://sourceforge.net/projects/bbmap/). We performed normalisation and assembly separately for each of the 167 samples. We then used the resulting scaffolds to search a database formed by combining the NCBI ‘refseq protein’ database with the viruses from NCBI ‘nr’ database. The search was performed using Diamond blastx (version 0.9.31; Buchfink *et al*. 2014) with an e-value threshold of 1×10^-30^, permitting frameshifts, and retaining hits within 5% of the top hit.

The resulting sequences were examined to exclude all phage, retroelements, giant viruses (i.e., Mimiviruses and relatives), and likely contaminants such as perfect matches to well-characterised plant, human, pet, and vertebrate livestock viruses (e.g. Ebola virus, Hepatitis B virus, Bovine viral diarrhoea virus, Murine leukemia virus). We also excluded virus fragments that co-occurred across samples with species other than *Drosophila*, such as mites and fungi, as likely to be viruses of those taxa. Our remaining candidate virus list included known and potentially novel DNA viruses, and one previously reported *Drosophila* RNA virus. For each of these viruses we selected at least one representative population sample, based on high coverage, for targeted genome re-assembly.

For targeted re-assembly of each virus we remapped all non-normalised reads to the putative virus scaffolds from the first assembly and retained all read pairs for which at least one partner had mapped. Using these virus-enriched read sets we then performed a second *de novo* SPAdes assembly for each target sample (as above), but to aid scaffolding and repeat resolution we additionally included the long reads (Antipov *et al*. 2015) that had been generated separately from UA_Yal_14_16, ES_Gim_15_30, UA_Ode_16_47 and UA_Kan_16_57. We examined the resulting assembly graphs using Bandage version 0.8.1 (Wick *et al*. 2015), and based on inspection of coverage and homology with related viruses we manually resolved short repeat regions, bubbles associated with polymorphism, and long terminal repeat regions. For viruses represented by very few low-coverage fragments, we concentrated assembly and manual curation on genes and gene fragments that would be informative for phylogenetic analysis.

For *Drosophila* Vesanto virus, a bidna-like virus with two previously-reported segments (Kapun *et al*. 2020), a preliminary manual examination of the assembly graph identified a potential third segment. We therefore took two approaches to explore the possibility that this virus is composed of more than two segments. First, to identify completely new segments, we mapped reads from samples with or without segments S01 and S02 to all high-coverage scaffolds from one sample that contained those segments. This allowed us to identify possible further segments based on their pattern of co-occurrence across samples (e.g. Batson *et al*. 2020, Obbard *et al*. 2020). Second, to identify substantially divergent (but homologous) alternative segments we used a blastp similarity search using predicted Vesanto virus proteins and predicted proteins from *de novo* scaffolds (e-value 10^-20^). Again, we examined targeted assembly graphs using Bandage (Wick *et al*. 2015), and resolved inverted terminal repeats and apparent mis-assemblies manually.

To annotate viral genomes with putative coding DNA sequences we used getORF from the EMBOSS package (Rice *et al*. 2000) to identify all open reading frames of 150 codons or more that started with ATG, and translated these to provide putative protein sequences. We retained those with substantial similarity to known proteins from other viruses, along with those that did not overlap longer open reading frames.

### Presence of DNA viruses in publicly available Drosophila datasets

To detect all known and novel *Drosophila* DNA viruses present in publicly available DNA *Drosophila* datasets, we chose 28 ‘projects’ from the NCBI Sequence Read Archive and mapped these to virus genomes using Bowtie 2 (Langmead and Salzberg 2012). Among these were several projects associated with the *Drosophila melanogaster* Genome Nexus (Lack *et al*. 2015, Lange *et al*. 2016, Sprengelmeyer *et al*. 2019), the Drosophila Real-Time Evolution Consortium (Dros-RTEC; Machado *et al*. 2019, Kapun *et al*. 2021), pooled GWAS studies (e.g. Endler *et al*. 2018), evolve-and-resequence studies (Jalvingh *et al*. 2014, Schou *et al*. 2017, Kelly and Hughes 2019), studies of local adaptation (e.g. Campo *et al*. 2013, Kang *et al*. 2019), and introgression (Kao *et al*. 2015). In total this represented 3003 Illumina sequencing ‘run’ datasets. The ‘project’ and ‘run’ identifiers are listed in figshare repository 10.6084/m9.figshare.14161250 file S7. For each run, we mapped up to 10 million reads to *Drosophila* DNA viruses (forward reads only for paired-end datasets) using Bowtie 2, and recorded the best-mapping location for each read, as above. Short reads and low complexity regions allow some cross-mapping among the larger viruses, and between viruses and the fly genome. We therefore chose an arbitrary detection threshold of 250 mapped reads to define the presence of each of the larger viruses (expected genome size >100 kbp) and a threshold of 25 reads for the smaller viruses (genome size <100 kbp). Consequently, our estimates may be conservative tests of virus presence, and the true prevalence may be slightly higher. We additionally selected a subset of the public datasets for *de novo* assemblies of Vesanto virus (datasets ERR705977, ERR173251, ERR2352541, SRR3939080), an Adintovirus (SRR3939056), and Galbut virus (SRR5762793, SRR1663569), using the same assembly approach as outlined for DrosEU data above.

### Phylogenetic inference

To infer the phylogenetic relationships among DNA viruses of *Drosophila* and representative viruses of other species, we selected a small number of highly conserved virus protein-coding loci that have previously been used for phylogenetic inference. For Densoviruses we used the viral replication initiator protein, NS1 (Pénzes *et al*. 2020), for Adintoviruses and Bidna-like viruses we used DNA Polymerase B (Krupovic and Koonin 2014, Starrett *et al*. 2020), for Poxviruses we used rap-94, and the large subunits of Poly-A polymerase and the mRNA capping enzyme (Thézé *et al*. 2013), and for Nudiviruses, Filamentous viruses and Hytrosaviruses we used P74, Pif-1, Pif-2, Pif-3, Pif-5 (ODV-e56) and the DNA polymerase B (e.g., Kawato *et al*. 2019). In each case we used a blastp search to identify a representative set of similar proteins in the NCBI ‘nr’ database, and among proteins translated from publically available transcriptome shotgun assemblies deposited in GenBank. For the Nudiviruses, Filamentous viruses and Hytrosaviruses we combined these with proteins collated by Kawato *et al* (2019). We aligned protein sequences for each locus using t-coffee mode ‘accurate’, which combines structural and profile information from related sequences (Notredame *et al*. 2000), and manually ‘trimmed’ poorly aligned regions from each end of each alignment. We did not filter the remaining alignment positions for coverage or alignment ‘quality’, as this tends to bias toward the guide tree and to give false confidence (Tan *et al*. 2015). We then inferred trees from concatenated loci (where multiple loci were available) using IQtree2 with default parameters (Minh *et al*. 2020), including automatic model selection and 1000 ultrafast bootstraps.

### Age of an endogenous viral element

To infer the age of an endogenous copy (EVE) of Galbut virus (a dsRNA Partitivirus Cross *et al*. 2020), we used a strict-clock Bayesian phylogenetic analysis of virus sequences, as implemented in BEAST 1.10.2 (Suchard *et al*. 2018). To make this inference our assumption is that any evolution of the EVE after insertion is negligible relative to RNA virus evolutionary rates. We assembled complete 1.6 kb segment sequences from publicly-available RNA sequencing datasets (Lin *et al*. 2016, Garlapow *et al*. 2017, Yablonovitch *et al*. 2017, Bost *et al*. 2018, Shi, White, *et al*. 2018, Everett *et al*. 2020), and filtered these to retain unique sequences and exclude possible recombinants identified with GARD (Kosakovsky Pond *et al*. 2006). The few recombinants were all found in multiply-infected pools, suggesting they may have been chimeric assemblies. For sequences from Shi et al. (2018) we constrained tip dates according to the extraction date, and for other studies we constrained tip dates to the three-year interval prior to project registration. We aligned these sequences with the EVE sequence, and during phylogenetic analysis we constrained most recent date for the EVE to be its extraction date, but left the earliest date effectively unconstrained. Because the range of virus tip dates covered less than 10 years we imposed time information through a strongly informative lognormal prior on the strict clock rate, chosen to reflect the spread of credible evolutionary rates for RNA viruses (e.g., Peck and Lauring 2018). Specifically, we applied a data-scale mean evolutionary rate of 4×10^-4^ events/site/year with standard deviation 2.5×10^-4^, placing 95% of the prior density between 1×10^-3^ and 1.3×10^-4^. As our sampling strategy was incompatible with either a coalescent or birth-death tree process, we used a Bayesian Skyline coalescent model to allow flexibility in the coalescence rate, and thereby minimise the impact of the tree prior on the date (although alternative models gave qualitatively similar outcomes). We used the SDR06 substitution model (Shapiro *et al*. 2006) and otherwise default priors, running the MCMC for 100 million steps and retaining every 10 thousandth state. The effective sample size was greater than 1400 for every parameter. BEAST input xml is provided via figshare repository 10.6084/m9.figshare.14161250.

### Virus quantification, and the geographic and temporal distribution of viruses

To quantify the (relative) amount of each virus in each pooled sample, we mapped read pairs that had not been mapped concordantly to the *Drosophila* microbiome reference (above) to the virus genomes. This approach means that low complexity reads map initially to the fly and microbiota, and are thus less likely to be counted or mis-mapped among viruses. This slightly reduces the detection sensitivity (and counts) but also increases the specificity. We mapped using Bowtie 2 (Langmead and Salzberg 2012), recording the best mapping location as above. We used either read count (per million reads) divided by target length (per kilobase) to quantify the viruses, or this value normalised by the equivalent number for *Drosophila* (combined *D. melanogaster* and *D. simulans* reads) to provide an estimate of virus genomes per fly genome in each pool. To quantify Vesanto virus genomes we excluded terminal inverted repeats from the reference, as these may be prone to cross-mapping among segments.

To provide a simple estimate of prevalence, we assumed that pools represented independent samples from a uniform global population, and assumed that a pool of *n* flies constituted *n* Bernoulli trials in which the presence of virus reads indicated at least one infected fly (e.g., Speybroeck *et al*. 2012). Based on this model, we inferred a maximum-likelihood estimate of global prevalence for each virus, with 2 log-likelihood intervals. Because some cross-mapping between viruses is possible, and because barcode switching can cause reads to be mis-assigned among pools, we chose to use a virus detection threshold of 1% of the fly genome copy number to define ‘presence’. This threshold was chosen on the basis that male flies artificially infected with Kallithea virus have a virus genome copy number 5-fold higher than that of the fly three days post infection (Palmer *et al*. 2018), or around 1% of the fly genome copy number for a single infected fly in a pool of 40. Thus, although our approach may underestimate virus prevalence if titre is low, it provides some robustness to barcode switching while also giving reasonable power to detect a single infected fly.

In reality, pools are not independent of each other in time or space, or other potential predictors of viral infection. Therefore, for the three most prevalent viruses (*Drosophila* Kallithea nudivirus, *Drosophila* Linvill Road densovirus, and *Drosophila* Vesanto virus) we analysed predictors of the presence and absence of each virus within population pools using a binomial generalised linear mixed model approach. We fitted linear mixed models in a spatial framework using R-INLA (Blangiardo *et al*. 2013), taking a Deviance Information Criterion (DIC) of 2 or larger as support for a spatial or spatiotemporal component in the model. In addition to any spatial random effects, we included one other random-effect and four fixed-effect predictors. The fixed effects were: the level of *D. simulans* contamination (measured as the percentage *D. simulans* Ago2 reads); the amount of *Wolbachia* (measured as reads mapping to *Wolbachia* as relative to the number mapped to fly genomes); the sampling season (early or late); and the year (unordered categorical 2014, 2015, 2016). We included sampling location as a random effect, to account for any additional non-independence between collections made at the same sites or by the same collector. The inclusion of a spatially distributed random effect was supported for *Drosophila* Kallithea nudivirus and *Drosophila* Linvill Road densovirus, but this did not vary significantly with year. Map figures were plotted and model outputs summarised with the R package ggregplot (https://github.com/gfalbery/ggregplot), and all code to perform these analyses is provided via figshare repository 10.6084/m9.figshare.14161250.

### Virus genetic diversity

Reads that had initially been mapped to *Drosophila* Kallithea nudivirus, *Drosophila* Linvill Road densovirus and *Drosophila* Vesanto virus (above) were remapped to reference virus genomes using BWA MEM with local alignment (Li 2013). For the segmented Drosophila Vesanto virus, we included multiple divergent haplotypes in the reference but excluded terminal inverted repeats, as reads derived from these regions will not map uniquely. After identifying the most common haplotype for each *Drosophila* Vesanto virus segment in each of the samples, we remapped reads to a single reference haplotype per sample. For all viruses, we then excluded secondary alignments, alignments with a Phred-scaled mapping quality (MAPQ) <30, and optical and PCR duplicates using picard v.2.22.8 ‘MarkDuplicates’ (http://broadinstitute.github.io/picard/). Finally, we excluded samples that had a read-depth of less than 25 across 95% of the mapped genome.

In addition to calculating per-sample diversity, to calculate total population genetic diversity we created single global pool, representative of diversity across the whole population, by merging sample bam files for each virus or segment haplotype. To reduce computational demands, each was down-sampled to an even coverage across the genome (no greater read depth at a site than the original median) and no sample contributed more than 500-fold coverage. To produce the final dataset for analyses, bam files for the global pool and each of the population pools were re-aligned around indels using GATK v3.8 (Van der Auwera *et al*. 2013). We created mPileup files using SAMtools (Li *et al*. 2009) to summarise each of these datasets using (minimum base quality = 40 and minimum MAPQ = 30), down-sampling population samples to a maximum read depth of 500. We masked regions surrounding indels using ‘po-poolation’ (Kofler, Orozco-terWengel, *et al*. 2011), and generated allelic counts for variant positions in each using ‘popoolation2’ (Kofler, Pandey, *et al*. 2011), limiting our search to single nucleotide polymorphisms (SNPs) with a minor allele frequency of at least 1%.

To calculate average pairwise nucleotide diversity at synonymous (TT_S_) and non-synonymous (TT_A_) sites we identified synonymous and non-synonymous SNPs using popoolation (Kofler, Orozco-terWengel, *et al*. 2011), excluding SNPs with a minor allele frequency of less than 1%. In general, estimates of genetic diversity from pooled samples, such as those made by popoolation and population2, attempt to account for variation caused by finite sample sizes of individuals each contributing to the pool of nucleic acid. However, such approaches cannot be applied to viruses from pooled samples, as it is not possible to infer the number of infected flies in the pool or even to equate an infected fly with an individual (flies may be multiply infected). For this reason, we calculated TT_A_ and TT_S_ based on raw allele counts derived from read frequencies (code is provided via figshare repository 10.6084/m9.figshare. 14161250). We did this separately for each gene in the merged global pool, and also for the whole genome in each infected population pool.

### Structural variation and indels in Drosophila Kallithea nudivirus

Large DNA viruses such as *Drosophila* Kallithea nudivirus can harbour transposable element (TE) insertions and structural rearrangements (Loiseau *et al*. 2020), and often contain abundant short repeat-length variation (Zhao *et al*. 2012). To identify large-scale rearrangements, we identified all read pairs for which at least one read mapped to *Drosophila* Kallithea nudivirus, and used SPAdes (Bankevich *et al*. 2012) to perform *de novo* assemblies separately for each dataset using both *in silico* normalised and un-normalised reads. We then selected those scaffolds approaching the expected length of the genome (>151 Kbp), and examined the assembly graphs manually using bandage (Wick *et al*. 2015), retaining those in which a single circular scaffold could be seen, with a preference for un-normalised datasets. These were then linearised starting at the DNA Polymerase B coding sequence, and aligned using muscle (Edgar 2004). This approach will miss structural variants at low frequency within each population, but could identify any major rearrangements that are fixed differently across populations.

To detect polymorphic transposable element insertions that were absent from the reference genome, we identified 16 population samples that had more than 300-fold read coverage of *Drosophila* Kallithea nudivirus and extracted all reads that mapped to the virus. We aligned these to 135 *D. melanogaster* TEs curated in the November 2016 version of Repbase (Bao *et al*. 2015) using blastn (-task megablast). All reads for which one portion aligned to the virus (Genome reference KX130344.1) and another portion aligned to a *D. melanogaster* TE were identified as chimeric using the R script provided by Peccoud *et al*. (2018), and those for which the read-pair spanned TE ends were considered evidence of a TE insertion.

Finally, to catalogue short indel polymorphisms in coding and intergenic regions, we used po-poolation2 (Kofler, Pandey, *et al*. 2011) to identify the genomic positions (relative to the reference genome) in each of the infected samples for which a gap is supported by at least 5 reads. We used a chi-square test for independence to test if there was an association between the coding status of a position and the probability that an indel was supported at that position in at least one population sample.

## Results

Over six and a half thousand flies were collected from 47 locations across Europe across three years as part of the DrosEU project (Kapun *et al*. 2020, Kapun *et al*. 2021). Their DNA was sequenced in population pools of around 40 flies, resulting in a total of 8.4 billion trimmed read pairs, with between 27.3 and 78 million pairs per sample. Using these reads we find evidence for 14 distinct DNA viruses associated with *Drosophila melanogaster* in Europe, of which nine have not been previously reported. We find two of the viruses to occur at a relatively high prevalence of 2-3%, but most are extremely rare.

### Host species composition

On average, 93% of reads (range 70 - 98%) could be mapped to *Drosophila* or likely components of the *Drosophila* microbial community. *Wolbachia* made up an average of 0.5% of mapped non-fly reads (range 0.0 - 2.9%); other mapped bacterial reads together were 0.6% (0.0 - 3.2%), and microbial eukaryotes were 0.3% (0.0 - 3.7%). The eukaryotic microbiota included the fungal pathogen *Entomophthora muscae* (e.g. Elya *et al*. 2018), with reads present in 42 of 167 samples (up to 1.38 reads per kilobase per million reads, RPKM), a novel trypanosomatid distantly related to *Herpetomonas muscarum* (e.g. Sloan *et al*. 2019) with reads present in 80 samples (up to 0.87 RPKM). We also identified the microsporidian *Tubulinosema ratisbonensis* (e.g. Niehus *et al*. 2012) in one sample (0.54 RPKM). We excluded two virus-like DNA Polymerase B fragments from the analyses because they consistently co-occurred with a fungus very closely related to *Candida* (*Clavispora*) *lusitaniae* (correlation coefficient on >0.94, *p*<10^-10^; figshare repository 10.6084/m9.figshare.14161250 File S4). For a detailed assessment of the microbial community in the 2014 collections, see Kapun *et al* (2020) and Wang *et al* (2020). Raw and normalised read counts are presented in figshare repository 10.6084/m9.figshare.14161250 File S3, and raw data are available from the Sequence Read Archive under project accession PRJNA388788.

The remaining 2% to 30% of reads could include metazoan species associated with *Drosophila*, such as nematodes, mites, or parasitoid wasps. By mapping all reads to small reference panel of *Cytochrome Oxidase I* (COI) sequences (figshare repository S2), we identified 13 samples with small read numbers mapping to potentially parasitic nematodes, including an unidentified species of *Steinernema*, two samples with reads mapping to *Heterorhabditis bacteriophora* and three with reads mapping to *Heterorhabditis marelatus*. *De novo* assembly also identified an 8.4 kbp nematode scaffold with 85% nucleotide identity to the mitochondrion of *Panagrellus redivivus*, a free-living rhabditid associated with decomposing plant material. Reads from this nematode were detectable in 73 of the 167 samples, rarely at a high level (up to 0.8 RPKM). Only one sample contained reads that mapped to mite COI, sample UK_Dai_16_23, which mapped at high levels (5.8 and 2.2 RPKM) to two unidentified species of Parasitidae (Mesostigmata, Acari). We excluded two Cyclovirus-like fragments from the analyses below because they occurred only in the sampled contaminated with the two mites, suggesting that they may be associated with the mites or integrated into their genomes (figshare repository 10.6084/m9.figshare.14161250 S3 and S4).

To detect the presence of drosophilid hosts other than *D. melanogaster*, we mapped all reads to a curated panel of short diagnostic sequences from COI and Argonaute-2, the latter chosen for its ability to reliably distinguish between the close relatives *D. melanogaster* and *D. simulans*. As expected from previous analyses of these data (e.g. Kapun *et al*. 2020), 30 of the 167 samples contained *D. simulans* at a threshold of >1% of Ago2 reads. Mapping to COI sequences from different species, we identified only three further *Drosophila* species present in any sample at a high level. These included two small yellowish European species; *D. testacea*, which accounted for 2.4% of COI in UA_Cho_15_26 (263 reads), and *D. phalerata*, which accounted for 12.2% of COI in AT_Mau_15_50 (566 reads). The presence of both species was confirmed by additional mapping to *Adh*, *Amyrel*, *Gpdh* and *6-PGD* (figshare repository 10.6084/m9.figshare.14161250 S3), and their mitochondrial genomes were recovered as 9 kbp and 16 kbp *de novo* scaffolds, respectively. More surprisingly, some of the collections made in 2015 contained reads derived from *D. serrata*, a well-studied species closely related to *D. melanogaster* and endemic to tropical Australia (Reddiex *et al*. 2018). Samples TR_Yes_15_7 and FR_Got_15_48 had particularly high levels of *D. serrata* COI, with 94% (23,911 reads) and 7% (839 reads) of COI respectively, but reads were also detectable in another 6 pools. The presence of *D. serrata* sequences was confirmed by mapping to *Adh*, *Amyrel*, *Gpdh* and *6-PGD* (figshare repository 10.6084/m9.figshare.14161250 S3). However, examination of splice junctions showed that *D. serrata* reads derived from cDNA rather than genomic DNA, and must therefore result from cross-contamination during sequencing or from barcode switching. Below, we note where conclusions may be affected by the presence of species of other than *D. melanogaster*.

Finally, among *de novo* assembled contigs, we also found evidence for several crop-plant chloroplasts and vertebrate mitochondria that are likely to represent sequencing or barcode-switching contaminants. The amounts were generally very low (median 0.01 RPKM), but a few samples stood out as containing potentially high levels of these contaminants. Most notably sample TR_Yes_15_7, in which only 76% of reads mapped to fly or expected microbiota, had 8.1 RPKM of human mtDNA, 5.1 RPKM of *Cucumis melo* cpDNA, and 3.5 RPKM of *Oryza sativa* cpDNA. We do not believe contamination of this sample has any impact on our findings.

### Previously-reported DNA virus genomes

Six different DNA viruses were previously detected among DrosEU samples from 2014 and reported by Kapun *et al*. (2020). These included one known virus (Drosophila Kallithea Nudivirus; Webster *et al*. 2015) and five new viruses, of which four were assembled by Kapun *et al*. (2020). *Drosophila* Kallithea nudivirus is a relatively common virus of *D. melanogaster* (Webster *et al*. 2015) that has a circular dsDNA genome of *ca*. 153 kbp encoding approximately 95 proteins (Figure 1), and is closely related to *Drosophila innubila* nudivirus (Figure 2A). *Drosophila* Esparto nudivirus is a second Nudivirus associated with *D. melanogaster* that was present at levels too low to permit assembly by Kapun *et al*. (2020), but was instead assembled in that paper from a *D. melanogaster* sample collected in Esparto, California USA (SRA dataset SRR3939042; Machado *et al*. 2019). It has a circular dsDNA genome of *ca*. 183 kbp that encodes approximately 90 proteins, and it is closely related to *Drosophila innubila* Nudivirus and *Drosophila* Kallithea nudivirus (Figure 1; Figure 2A). *Drosophila* Viltain densovirus and *Drosophila* Linvill Road densovirus are both small viruses related to members of the Parvoviridae, with ssDNA genomes of approximately 5 kb. *Drosophila* Viltain densovirus is most closely related to *Culex pipiens* ambidensovirus (Jousset *et al*. 2000), and the genome appears to encode at least four proteins—two in each orientation (Figure 1; Figure 2B). As expected, the ends of the genome are formed of short inverted terminal repeats (Figure 1). *Drosophila* Linvill Road densovirus is most closely related to the unclassified *Haemotobia irritans* densovirus (Ribeiro *et al*. 2019) and appears to encode at least three proteins, all in the same orientation (Figure 1; Figure 2B). As with *Drosophila* Esparto nudivirus, Kapun *et al*. (2020) were unable to assemble the *Drosophila* Linvill Road densovirus genome from the DrosEU 2014 data and instead based their assembly on a collection of *D. simulans* from Linvilla, Pennsylvania USA (SRR2396966; Machado *et al*. 2019). Here we identified a DrosEU 2016 collection (ES_Ben_16_32; Benalua, Spain) with sufficiently high titre to permit an improved genome assembly (submitted to GenBank under accession MT490308). This is 99% identical to the previous *Drosophila* Linvill Road densovirus assembly, but by examination of the assembly graph we were able to complete more of the inverted terminal repeats and extend the genome length to 5.4 kb (Figure 1). Table 1 provides a summary of all DNA viruses detectable in DrosEU data.

**Figure 1:**
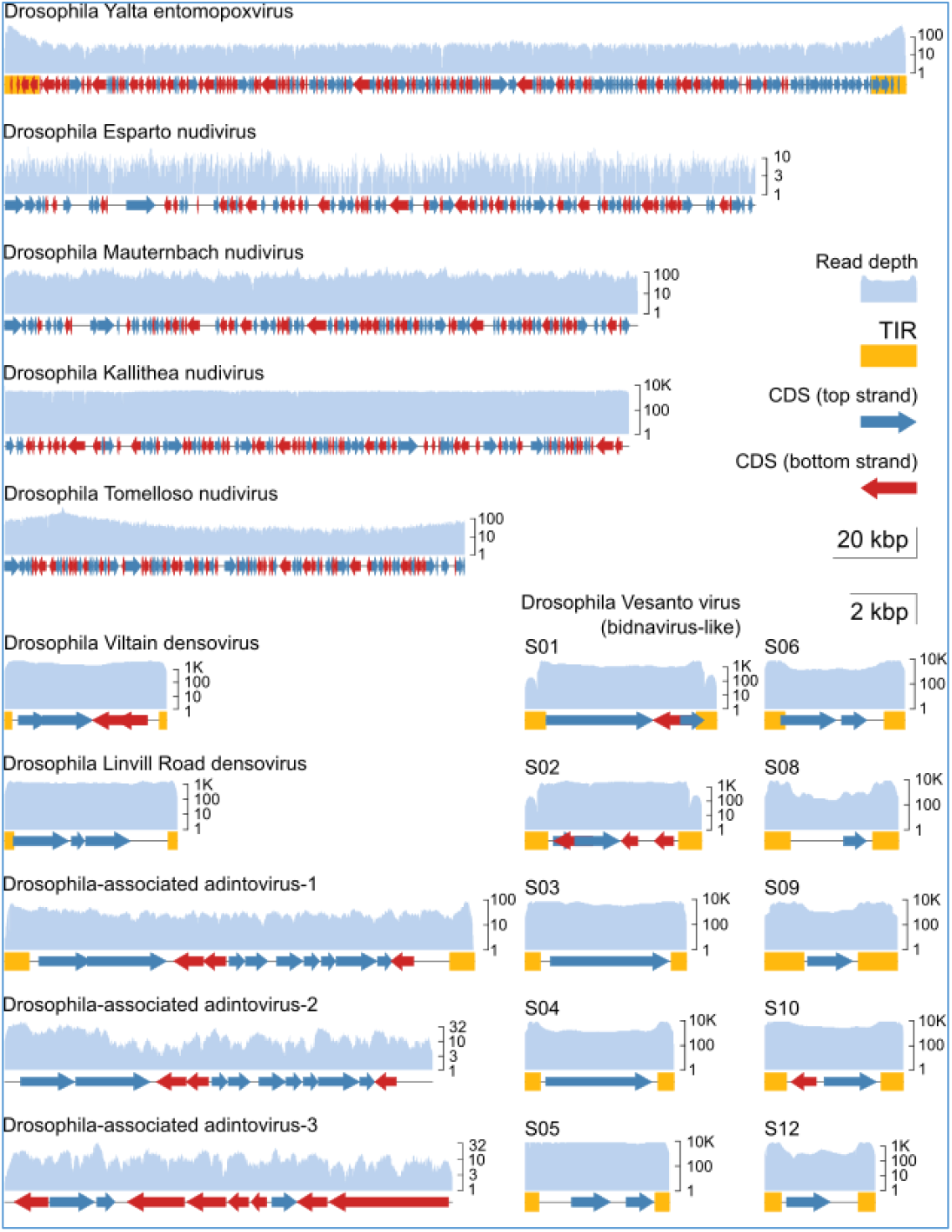
Genome structures and read depth. The plots show annotated coding DNA sequences (CDS, red and blue arrows), and terminal Inverted repeat (yellow boxes) for each of the near-complete virus genomes discussed. The read depth (pale blue) is plotted above the genome on a log scale for the population with the highest coverage in the DrosEU dataset. The five largest viruses (top) are plotted according to the 20 kbp scale bar, and the other viruses (bottom) are plotted according to the 2 kbp scale bar. The Nudiviruses are circular, and have been arbitrarily linearized for plotting. *Drosophila* Esparto nudivirus was completed using public dataset (SRR3939042). Note that *Drosophila* Vesanto virus segments S07 and S11 were absent from the illustrated sample (lower right).

**Figure 2:**
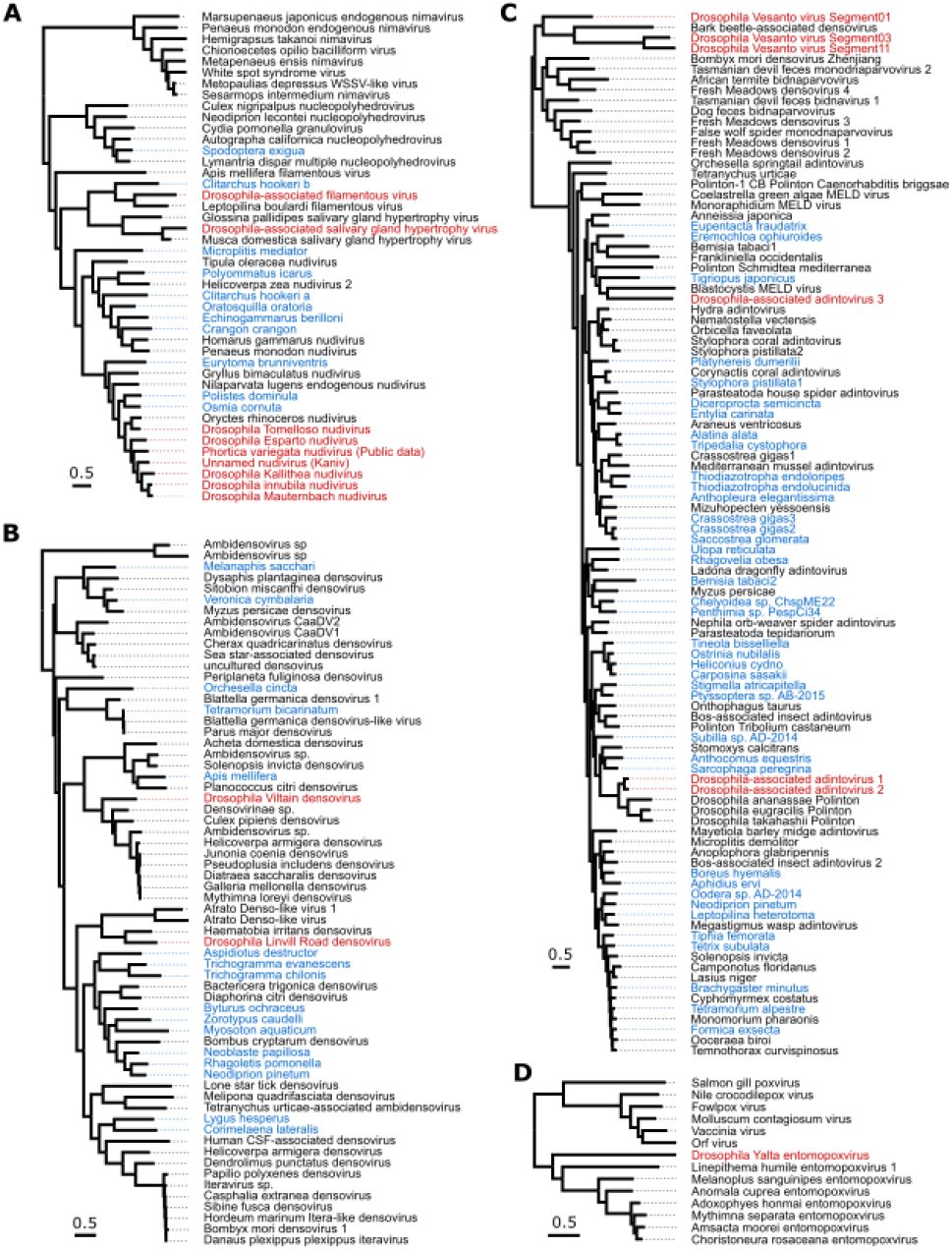
Phylogenetic relationships. (A) Nudiviruses, Hytrosaviruses, Filamentous viruses, Nucleopolyhedrosis viruses and Nimaviruses, inferred from six concatenated protein coding genes. Note that these lineages are extremely divergent, and the alignment is not reliable at deeper levels of divergence. (B) Densoviruses, inferred from NS1. (C) Bidnaviruses (sometimes labelled ‘Densovirus’) and Adintoviruses (including representative Polintons), inferred from DNA Polymerase B. (D) Pox and Entomopox viruses, inferred from three concatenated protein coding genes. All phylogenies were inferred from protein sequences by maximum likelihood, and scale bars represent 0.5 amino-acid substitutions per site. In each case, trees are mid-point rooted, viruses reported from *Drosophila* are shown in red, and sequences identified from virus transcripts in publicly-available transcriptome assemblies are shown in blue, labelled by host species. The Nudivirus from *Phortica variegata* was derived from PRJNA196337 (Vicoso and Bachtrog 2013). Alignments and tree files with bootstrap support are available through FisgShare repository.

**Table 1:** DNA Viruses of *Drosophila* present in the DrosEU dataset

| Virus name | Relationship | Genome status | Assembly (bp) | First published | Genbank accessions | DrosEU code | Genome collection location | Source data |
| --- | --- | --- | --- | --- | --- | --- | --- | --- |
| *Drosophila* Kallithea nudivirus | Nudi-viruses | Complete | 152,388 | Webster et al. (2015) | KX130344 | UA_Kha_14_46 | Kharkiv, Ukraine | SRR5647730 |
| *Drosophila* Esparto nudivirus | Nudi-viruses | Complete | 183,261 | Kapun et al. (2020) | KY608910 | (Dros-RTEC) | Esparto, California | SRR3939042 |
| *Drosophila* Viltain densovirus | Den-sovi-ruses | Complete | 5,025 | Kapun et al. (2020) | KX648535 | FR_Vil_14_07 | Viltain, France | SRR5647729 |
| *Drosophila* Linvill Road densovirus | Den-sovi-ruses | Complete | 5,360 | Kapun et al. (2020)<br>Expanded here | MT490308 | ES_Ben_16_32 | Benalua, Spain | SRR8494475 |
| *Drosophila* Vesanto virus | Bidna-like vi-ruses | Up to 12 Seg-ments | Up to 52,097 | Kapun et al. (2020)<br>Expanded here | MT496850-MT496878 | UA_Kan_16_57<br>UA_Dro_16_56<br>FR_Got_15_48 | Kaniv, Ukraine Drogo-bych, Ukraine<br>Gotheron, France | SRR8494448<br>SRR8494441<br>SRR8439123 |
| *Drosophila* Tomelloso nudivirus | Nudi-viruses | Complete | 112,307 | This Paper | KY457233 | ES_Tom_15_28 | Tomelloso, Spain | SRR8439136 |
| *Drosophila* Mauternbach nudivirus | Nudi-viruses | Complete | 154,465 | This Paper | MG969167 | AT_Mau_15_50 | Mauternbach, Austria | SRR8439127 |
| *Drosophila* Yalta entomopoxvirus | Ento-mopox viruses | Complete | 219,929 | This Paper | MT364305 | UA_Yal_14_16 | Yalta, Ukraine | SRR5647764 |
| *Drosophila*-associated nudivirus (Kaniv) | Nudi-viruses | Fragmentary | 4,503 | This Paper | MT496841-MT496846 | UA_Kan_16_57 | Kaniv, Ukraine | SRR8494448 |
| *Drosophila*-associated filamentous virus | Fila-men-tous viruses | Fragmentary | 86,478 | This Paper | MT496832-MT496840 | ES_Gim_15_30 | Gimenells, Spain | SRR8439138 |
| *Drosophila*-associated hytrosavirus | Hy-trosa-viruses | Fragmentary | 16,606 | This Paper | MT469997-MT470014 | UA_Ode_16_47 | Odesa, Ukraine | SRR8494427 |
| *Drosophila*-associated adintovirus 1 | Adinto-like vi-ruses | Complete | 14,567 | This Paper | MT496847 | UA_Cho_15_26 | Kopachi (Chornobyl Ex-clusion Zone), Ukraine | SRR8439134 |
| *Drosophila*-associated adintovirus 2 | Adinto-like viruses | Near-Complete | 13,277 | This Paper | MT496848 | AT_Mau_15_50 | Mauternbach, Austria | SRR8439127 |
| *Drosophila*-associated adintovirus 3 | Adinto-like viruses | Near-Complete | 13,883 | This Paper | MT496849 | DK_Kar_16_4 | Karensminde, Denmark | SRR8494437 |

### Drosophila Vesanto virus may be a multi-segmented bidna-like virus

Kapun *et al*. (2020) also reported two segments of a putative ssDNA Bidnavirus, there called ‘Vesanto virus’ for its collection site in 2014 (submitted to GenBank in 2016 as KX648533 and KX648534). This was presumed to be a complete genome based on homology with *Bombyx mori* bidensovirus (Li *et al*. 2019). Here we have been able to utilise expanded sampling and a small number of long-read sequences to extend these segments and to identify multiple co-occurring segments.

While examining an assembly graph of sample UA_Kan_16_57, we noted a third scaffold with a similarly high coverage (>300-fold) and structure (4.8 kb in length with inverted terminal repeats). This sequence also appeared to encode a protein with distant homology to *Bidnavirus* DNA polymerase B, and we reasoned that it might represent an additional virus. We therefore mapped reads from datasets that had high coverage of *Drosophila* Vesanto virus segments S01 and S02 to all scaffolds from the *de novo* build of UA_Kan_16_57, with the objective of finding any additional segments based on their co-occurrence across datasets (e.g. as done by Batson *et al*. 2020, Obbard *et al*. 2020). This identified several possible segments, all between 3.3 and 5.8 kbp in length and possessing inverted terminal repeats. We then used their translated open reading frames to search all of our *de novo* builds, and in this way identified a total of 12 distinct segments that show structural similarity and a strong pattern of co-occurrence (Figure 1 and Figure 3; figshare repository 10.6084/m9.figshare.14161250 S5). To capture the diversity present among these putative viruses, we made targeted *de novo* builds of three datasets, incorporating both Illumina reads and Oxford nanopore reads (Table 1). We have submitted these contigs to GenBank as MT496850-MT496878, and additional sequences are provided in figshare repository 10.6084/m9.figshare.14161250 S6. Because the inverted terminal repeats and pooled sequencing of multiple infections make such an assembly particularly challenging, we also sought to support these structures by identifying individual corroborating Nanopore reads of 2 kbp or more. We believe the inverted terminal repeat sequences should be treated with caution, but it is nevertheless striking that many of these putative segments show sequence similarity in their terminal inverted repeats, as commonly seen for segmented viruses.

**Figure 3:**
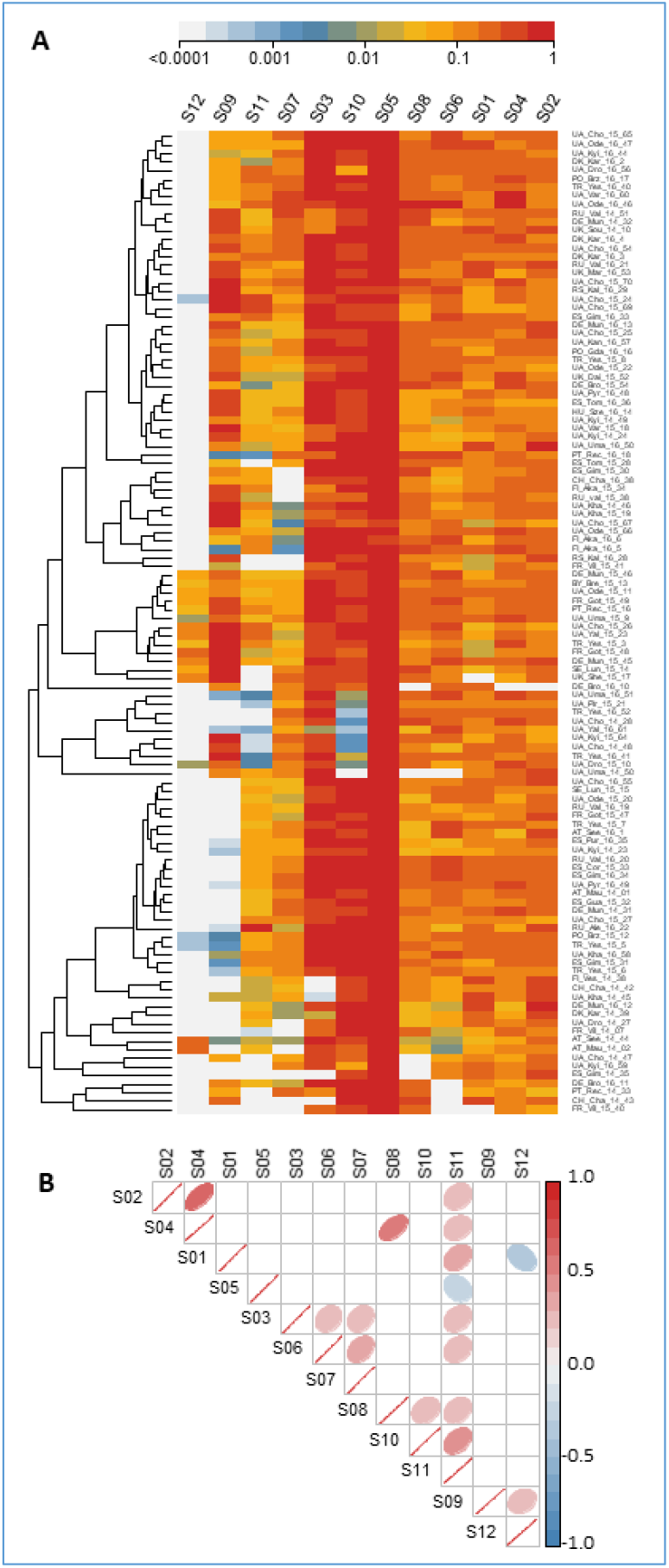
*Drosophila* Vesanto virus segment copy-number. (A) Heatmap showing the relative number of sequencing reads from each of the 12 Vesanto virus segments (columns), for each of the population samples (rows). Populations are included if at least one segment appeared at 1% of the fly genome copynumber. Rows and columns have been ordered by similarity (dendrogram) to identify structure within the data. Colours show copy-number relative to the high-est-copy segment, on a log scale. (B) Correlations in copy-number among the segments, with ‘significant’ correlations (p<0.05, no corrections) shown with coloured ellipses, according to the direction (red positive, blue negative) and strength of correlation. The absence of strong negative correlations between segments encoding homologous proteins (e.g. S01, S03, S11, which all encode genes with homology to DNA Polymerase B) may indicate that these segments do not substitute for each other.

**Figure 4:**
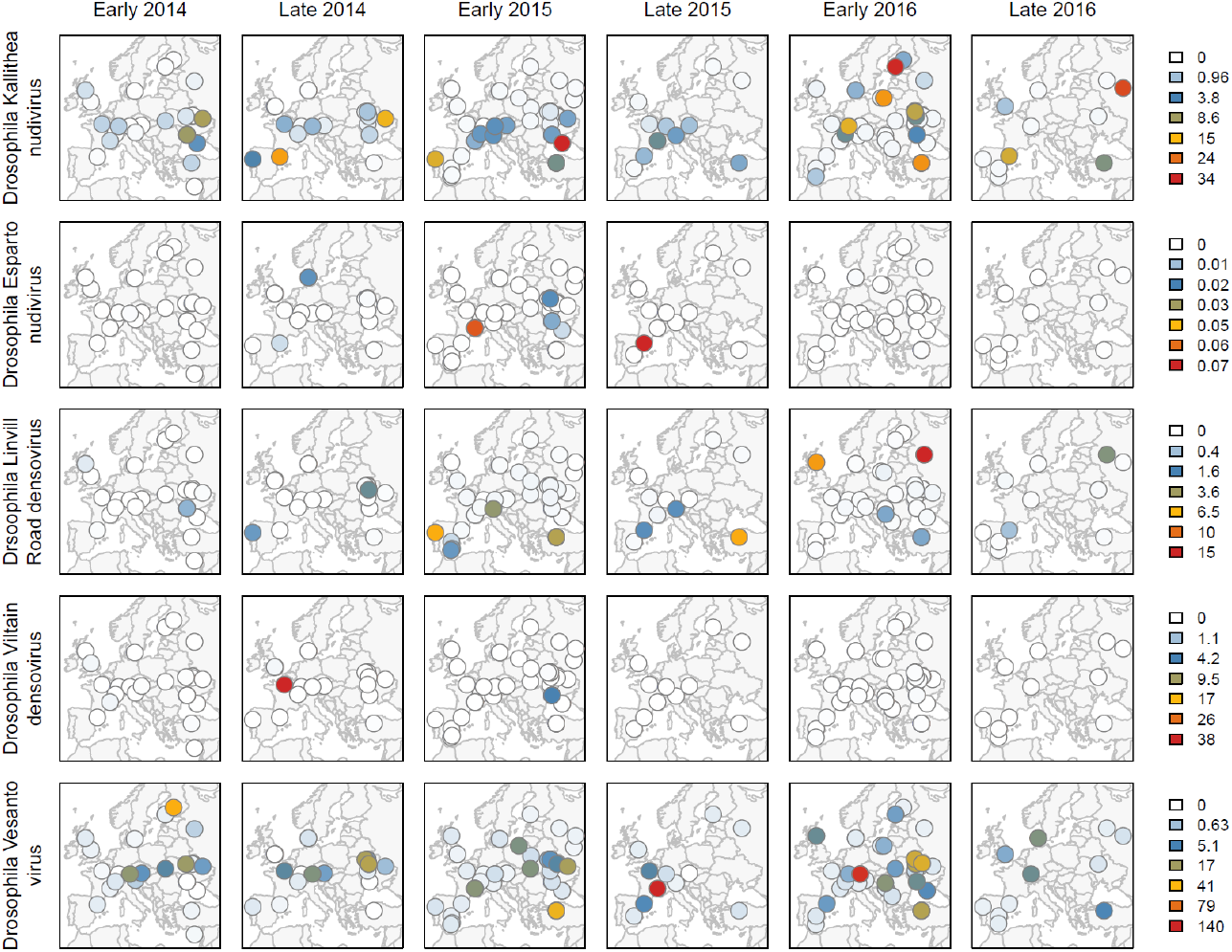
Geographic distribution of DNA virus reads in European *D. melanogaster*. Maps show the spatial distribution of virus read copy-number (relative to fly genomes) on a non-linear colour scale. Data are shown for the five viruses that were detected more than once (rows), separated by year and whether flies were collected relatively ‘early’ or ‘late’ in the season (columns).

Although we identified 12 distinct segments with strongly correlated presence/absence, not all segments were detectable in all affected samples (Figure 3A). Only segment S05, which encodes a putative glycoprotein and a putative nuclease domain protein, was always detectable in samples containing *Drosophila* Vesanto virus (in 91 of the 167 samples; figshare repository 10.6084/m9.figshare.14161250 S5). Several segments were very commonly detectable, such as S03 (protein with homology to DNA PolB) and S10 (encoding a protein with domain of unknown function DUF3472 and a putative glycoprotein) in around 70 samples, and segments S01, S02, S04, S06 and S08 in around 55 samples. Others were extremely rare, such as S12 (encoding a putative NACHT domain protein with homology to S09), which was only seen in five samples. We considered three possible explanations for this pattern.

Our first hypothesis was that *Drosophila* Vesanto virus has 12 segments, but that variable copy number among the segments causes some to occasionally drop below the detection threshold. In support of this, all segments are indeed detectable in the sample with the highest *Drosophila* Vesanto virus read numbers (FR_Got_15_49), ranging from 7-fold higher than the fly genome for S07 to 137-fold higher for S05. In addition, ‘universal’ segment S05 is not only the most widely-detected segment across samples, but also has the highest average read depth within samples. However, despite 1.6 million *Drosophila* Vesanto virus reads in the second highest copy-number sample (RU_Val_16_20; 125-fold more copies of S6 than of *Drosophila*), no reads mapped to S12, strongly suggesting the absence of S12 from this sample. Our second hypothesis was that some segments are ‘optional’ or satellite segments, or may represent alternative versions of other, homologous segments, comprising a reassorting community (as in influenza viruses). The latter is consistent with the apparent homology between some segments. For example, S01, S03, and S11 all encode DNA Polymerase B-homologs, and S06, S07 and S10 all encode DUF3472 proteins. It is also consistent with the universal presence of S05, which appears to lack homologs. However, two of the DNA PolB homologs are highly divergent (Figure 2C) to the extent it is hard to be confident of polymerase function, and we could not detect compelling negative correlations between homologous segments that might suggest that they substitute for each other in different populations (Figure 3B). Our third hypothesis was that ‘*Drosophila* Vesanto virus’ in fact represents multiple independent viruses (or phage), and that the superficially clear pattern of co-occurrence is driven by high (hypothetical) prevalence of this virus community in an occasional member of the *Drosophila* microbiota, such as a fungus or trypanosomatid. However, we were unable to detect any correlation with the mapped microbiota reads, and high levels of *Drosophila* Vesanto virus are seen in samples with few un-attributable reads. For example, sample PO_Brz_15_12 has 11-fold more copies of S6 than of the fly gneome, but less than 2% of reads derive from an unknown source (figshare repository 10.6084/m9.figshare.14161250 S3).

### The complete genome of a new divergent Entomopox virus

Kapun *et al*. (2020) also reported the presence of a pox-like virus in DrosEU data from 2014, but were unable to assemble the genome. By incorporating a small number of long sequencing reads, and using targeted reassembly combined with manual examination of the assembly graph, we were able to assemble this genome from dataset UA_Yal_14_16 (SRR5647764) into a single contig of 219.9 kb. As expected for pox-like viruses, the genome appears to be linear with long inverted terminal repeats of 8.4 kb, and outside of the inverted terminal repeats sequencing coverage was 15.7-fold (Figure 1). We suggest the provisional name ‘*Drosophila* Yalta entomopoxvirus’, reflecting the collection location (Yalta, Ukraine), and we have submitted the sequence to Genbank under accession number MT364305. This virus has very recently been shown to be most closely related to *Diachasmimorpha longicaudata* entomopoxvirus (Coffman and Burke 2020).

Within the *Drosophila* Yalta entomopoxvirus genome we identified a total of 177 predicted proteins, including 46 of the 49 core poxvirus genes, and missing only the E6R virion protein, the D4R uracil-DNA glycosylase, and the 35 kDa RNA polymerase subunit A29L (Upton *et al*. 2003). Interestingly, the genome has a higher GC content than the other previously published Entomopox viruses, which as a group consistently display the lowest GC content (< 21%) of the Poxvirus family (Perera *et al*. 2010, Thézé *et al*. 2013). Consistent with this, our phylogenetic analysis of three concatenated protein sequences suggests that the virus is distantly related, falling only slightly closer to Entomopox viruses than other pox viruses (Figure 2D). Given that all pox-like viruses infect metazoa, and that no animal species other than *D. melanogaster* appeared to be present in the sample, we believe *D. melanogaster* is likely to be the host.

### Two new complete Nudivirus genomes, and evidence for a third

In addition to *Drosophila* Kallithea nudivirus and *Drosophila* Esparto nudivirus, our expanded analysis identified three novel Nudiviruses that were absent from data collected in 2014. We were able to assemble two of these into complete circular genomes of 112.3 kb (27-fold coverage) and 154.5 kb (41-fold coverage), respectively, based on datasets from Tomelloso, Spain (ES_Tom_15_28; SRR8439136) and Mauternbach, Austria (AT_Mau_15_50; SRR8439127). We suggest the provisional names ‘*Drosophila* Tomelloso nudivirus’ and ‘*Drosophila* Mauternbach nudivirus’ for these viruses, reflecting the collection locations, and we have submitted the sequences to GenBank under accession numbers KY457233 and MG969167. We predict *Drosophila* Tomelloso nudivirus to encode 133 proteins (Figure 1), and phylogenetic analysis suggests that it is more closely related to a beetle virus (Oryctes rhinocerous Nudivirus, Figure 2A; Etebari *et al*. 2020) than to the other Nudiviruses described from *Drosophila*. *Drosophila* Mauternbach nudivirus is predicted to encode 95 proteins (Figure 1), and is very closely related to *Drosophila innubila* nudivirus (Figure 2A; Unckless 2011, Hill and Unckless 2018). However, synonymous divergence (*K*_S_) between these two viruses is approximately 0.7, i.e. nearly six-fold more than that between *D. melanogaster* and *D. simulans*, supporting their consideration as distinct ‘species’. The third novel Nudivirus was present at a very low level in a sample from Kaniv, Ukraine (UA_Kan_16_57, SRR8494448), and only small fragments of the virus could be assembled for phylogenetic analysis (Genbank accession MT496841-MT496846). This showed that the fragmentary nudivirus from Kaniv is approximately equally divergent from *D. innubila* nudivirus and *Drosophila* Mauternbach nudivirus (Figure 2A).

The collections from Tomelloso and Kaniv did not contain reads mapping to *Drosophila* species other than *D. melanogaster*, or to nematode worms or mites. Moreover, we identified *Drosophila* Tomelloso nudivirus in a number of experimental laboratory datasets from *D. melanogaster* (see below; Riddiford *et al*. 2020), and these lacked a substantial microbiome. Together these observations strongly support *D. melanogaster* as a host for these viruses. In contrast, COI reads suggest that the sample from Mauternbach may have contained one *Drosophila phalerata* individual (2.4% of diagnostic nuclear reads; figshare repository 10.6084/m9.figshare.14161250 S3). And, as we could not detect *Drosophila* Mauternbach nudivirus in any of the public datasets we examined (below), we reamain uncertain whether *D. melanogaster* or *D. phalerata* was the true host.

### Evidence for a new Filamentous virus and a new Hytrosavirus

Our search also identified fragments of two further large dsDNA viruses from lineages that have not previously been reported to naturally infect Drosophilidae. First, in sample UA_Ode_16_47 (SRR8494427) from Odesa, Ukraine, we identified around 16.6 kb of a novel virus related to the salivary gland hypertrophy viruses of *Musca domestica* and *Glossina palpides* (Figure 2A; Prompiboon *et al*. 2010, Kariithi *et al*. 2013). Our assembled fragments comprised 18 short contigs of only 1 to 3-fold coverage (submitted under accessions MT469997-MT470014). As the *Glossina* and *Musca* viruses have circular dsDNA genomes of 124.3 kbp and 190.2 kbp respectively, we believe that we have likely sequenced 5-15% of the genome. Because this population sample contains a small number of reads from *D. simulans* and an unknown nematode worm related to *Panagrellus redivivus*, and because we were unable to detect this virus in public datasets from *D. melanogaster* (below), the true host remains uncertain. However, given that the closest relatives all infect Diptera, it seems likely that either *D. melanogaster* or *D. simulans* is the host.

Second, in sample ES_Gim_15_30 (SRR8439138) from Gimenells, Spain, we identified around 86.5 kb of a novel virus distantly related to the filamentous virus of *Leptopilina boulardi*, a parasitoid wasp that commonly attacks *Drosophila* (Figure 2; Lepetit *et al*. 2016). The assembled fragments comprised 9 scaffolds of 5.9-16.9 kbp in length and 3 to 10-fold coverage, and are predicted to encode 69 proteins (scaffolds submitted to Genbank under accessions MT496832-MT496840). *Leptopilina boulardi* filamentous virus has a circular genome of 111.5 kbp predicted to encode 108 proteins. This suggests that, although fragmentary, our assembly may represent most of the virus. A small number of reads from ES_Gim_15_30 mapped to a relative of nematode *Panagrellus redivivus* and, surprisingly, to the Atlantic salmon (*Salmo salar*), but we consider these unlikely hosts as the level of contamination was very low and other filamentous viruses are known to infect insects. We were unable to detect the novel filamentous virus in any public datasets from *D. melanogaster* (below), and given that *Leptopilina boulardi* filamentous virus infects a parasitoid of *Drosophila*, it is possible that this virus may similarly infect a parasitoid wasp rather than the fly. However, as we were unable to detect any reads mapping to *Leptopilina* or other parasitoids of *Drosophila* in any of our samples, we think *D. melanogaster* is a good candidate to be a true host.

### Near-complete genomes of three Adintoviruses

Based on the presence of a capsid protein, it is thought that some Polinton-like transposable elements (also known as Mavericks) are actually horizontally-transmitted viruses (Yutin *et al*. 2015). Some of these have recently been proposed as the Adintoviridae, a family of dsDNA viruses related to Bidnaviridae and other PolB-encoding DNA viruses (Starrett *et al*. 2020). We identified three possible Adintoviruses in DrosEU data. The first, which we refer to as *Drosophila*-associated adintovirus 1, occurred in sample UA_Cho_15_26 (SRR8439134) from Kopachi (Chornobyl Exclusion Zone), Ukraine and comprised a single contig of 14.5 kb predicted to encode 12 proteins. Among these proteins are not only a DNA Polymerase B and an integrase, but also homologs of the putative capsid, virion-maturation protease, and FtsK proteins of Adintoviruses (Starrett *et al*. 2020), and possibly very distant homologs of Hytrosavirus gene MdSGHV056 and Ichnovirus gene AsIV-cont00038 (Figure 1). The second, which we refer to as *Drosophila*-associated adintovirus 2, is represented by a 13.3 kb contig assembled using AT_Mau_15_50 from Mauternbach, Austria (SRR8439127). It is very closely related to the first Adintovirus, and encodes an almost-identical complement of proteins (Figure 1). In a phylogenetic analysis of DNA PolB sequences, both fall close to sequences annotated as Polintons in other species of *Drosophila* (Figure 2C). It is notable that these two datasets are those that are contaminated by *D. testacea* (1.3%, 1 fly) and *D. phalerata* (2.4%, 1 fly), respectively. We therefore think it likely that *Drosophila*-associated adintovirus 1 and 2 are associated with those two species rather than *D. melanogaster*, and may potentially be integrated into their genomes. These sequences have been submitted to Genbank under accessions MT496847 and MT496848.

In contrast, *Drosophila*-associated adintovirus 3 was assembled using sample DK_Kar_16_4 from Karensminde, Denmark (SRR8494437), from which other members of the Drosophilidae were absent. It is similarly 13.8 kb long, and our phylogenetic analysis of DNA PolB places it within the published diversity of insect Adintoviruses—although divergent from other Adintoviruses or Polintons of *Drosophila* (Figure 2C; see also Starrett *et al*. 2020). However, this sequence is only predicted to encode 10 proteins and these are generally more divergent, perhaps suggesting that this virus is associated with a completely different host species, such as the nematode related to *Panagrellus redivivus* or a trypanosomatid—although these species were present at very low levels. The sequence has been submitted to Genbank under accession MT496849

### Prevalence varies among viruses, and in space and time

Based on a detection threshold of 1% of the *Drosophila* genome copy-number, only five of the viruses (*Drosophila* Kallithea nudivirus, *Drosophila* Vesanto virus, *Drosophila* Linvill Road densovirus, *Drosophila* Viltain densovirus and *Drosophila* Esparto nudivirus) were detectable in multiple population pools. The other nine viruses were each detectable in a single pool. For viruses in a single pool, a simple maximum-likelihood estimate of prevalence—assuming independence of flies and pools—is 0.015% (with an upper 2-Log likelihood bound of 0.07%). Among the intermediate-prevalence viruses, *Drosophila* Esparto nudivirus and *Drosophila* Viltain virus were detected in 5 pools each, corresponding to a prevalence of 0.08% (0.03-0.17%), and *Drosophila* Linvill road densovirus was detected in 21 pools, indicating a prevalence of 0.34% (0.21-0.51%). The two most common viruses were *Drosophila* Kallithea nudivirus, which was detected in 93 pools giving a prevalence estimate of 2.1% (1.6-2.5%), and *Drosophila* Vesanto virus, which was detected in 114 pools giving a prevalence estimate of 2.9% (2.4-3.5%). However, it should be noted that all flies were male, and if virus prevalence differs between males and females then these estimates could be misleading. Both virulence and titre are known to differ between the sexes (e.g. in Drosophila Kallithea nudivirus; Palmer *et al*. 2018), although differences in prevalence were not found for RNA viruses of *Drosophila* (Webster *et al*. 2015).

*Drosophila* Kallithea nudivirus, *Drosophila* Vesanto virus, and *Drosophila* Linvill Road densovirus were sufficiently prevalent to analyse their presence / absence across populations using a Bayesian spatial Generalised Linear Mixed Model. Our analysis identified a spatial component to the distribution of both *Drosophila* Kallithea nudivirus and *Drosophila* Linvill Road densovirus that did not differ significantly between years, with a higher prevalence of *Drosophila* Kallithea nudivirus in southern and central Europe, and a higher prevalence of *Drosophila* Linvill Road densovirus in Iberia (Figure 5A and B; ΔDIC of −13.6 and −17.2, respectively, explaining 15.5% and 32.8% of the variance). In contrast, *Drosophila* Vesanto virus showed no detectable spatial variation in prevalence, but did vary significantly over time, with a significantly lower prevalence in 2014 compared to the other years (2015 and 2016 were higher by 1.27 [0.42,2.16] and 1.43 [0.50,2.14] respectively). The probability of observing a virus did not depend on the sampling season or the amount of *Wolbachia* in the sample.

**Figure 5:**
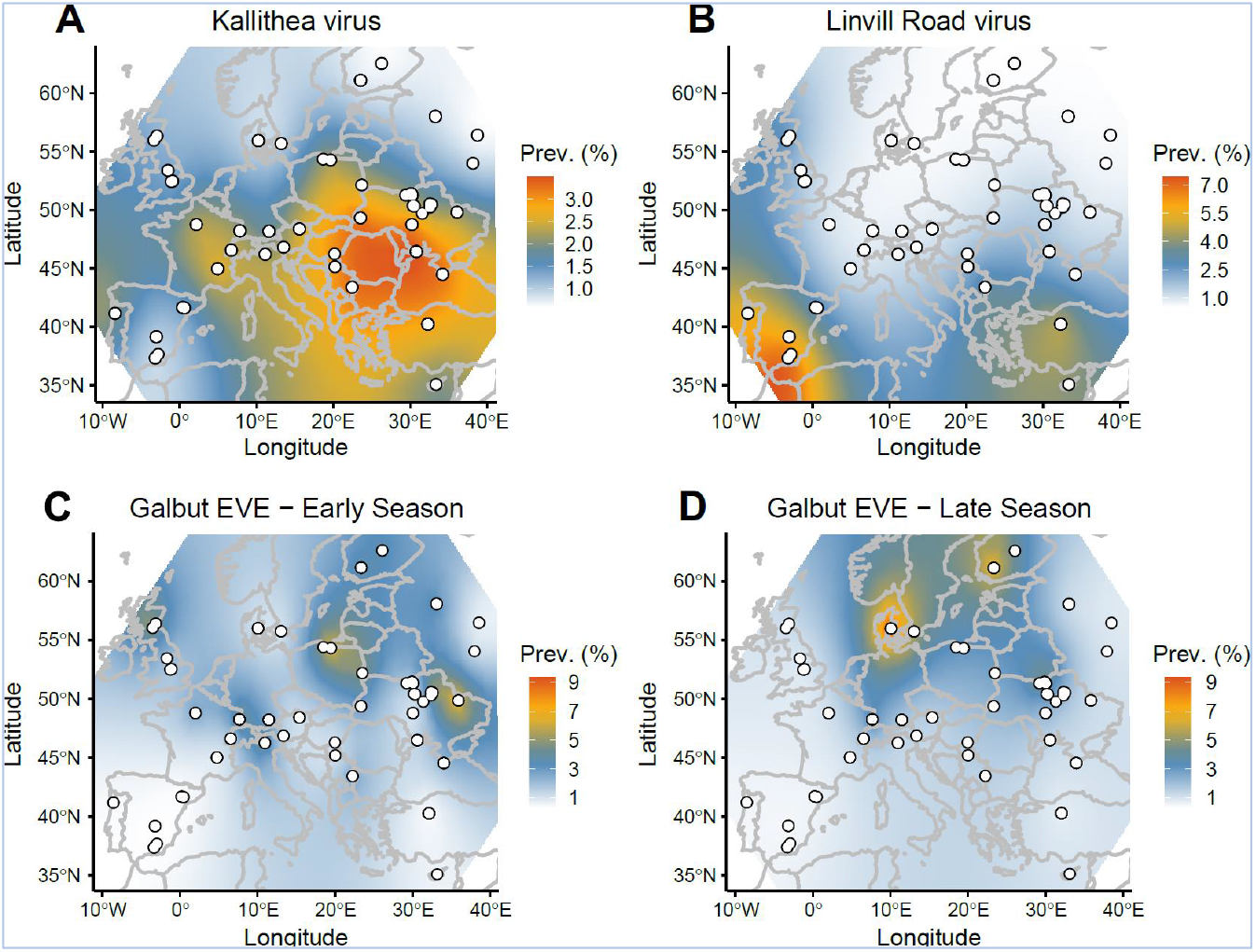
Geographic variation in estimated prevalence: *Drosophila* Kallithea nudivirus (A), *Drosophila* Linvill Road denosovirus (B), and the Galbut virus EVE (C and D). Sampling sites are marked as white dots, and the colour gradient illustrates predictions from the INLA model, but with scale transformed to the predicted individual-level prevalence (%), assuming independence among individuals and population samples of size 40. Only *Drosophila* Kallithea nudi virus, *Drosophila* Linvill road densovirus, and the Galbut virus EVE displayed a significant spatial component, and only the EVE differed between seasons.

As sampling location did not explain any significant variation in the probability of detecting any virus, it appears that—beyond broad geographic trends—there is little temporal consistency in virus prevalence at the small scale. At the broader geographic scale, it seems likely that climatic factors, directly or indirectly, play a role. For example, it may be that temperature and humidity affect virus transmission, as seen for many human viruses (Moriyama *et al*. 2020). Equally, host density and demography are strongly affected by climate, and will affect the opportunity for transmission, both within and between host species. For example, the probability of detecting *Drosophila* Linvill Road densovirus was positively correlated with the level of *D. simulans* contamination (95% credible interval of the log-odds ratio [2.9,14.6]), suggesting either that some reads derived from infections of *D. simulans* (in which the virus can have very high prevalence, see data from Signor *et al*. 2017), or that infections in *D. melanogaster* may be associated with spill-over from *D. simulans*.

### DNA viruses are detectable in publicly available Drosophila datasets

We wished to corroborate our claim that these viruses are associated with *Drosophila* by exploring their prevalence in laboratory populations and publicly available data. We therefore examined the first 10 million reads from each of 3003 sequencing runs from 28 *D. melanogaster* and *D. simulans* sequencing projects. In general, our survey suggests that studies using isofemale or inbred laboratory lines tend to lack DNA viruses (e.g., Mackay *et al*. 2012, Grenier *et al*. 2015, Lack *et al*. 2015, Gilks *et al*. 2016, Lange *et al*. 2016). In contrast, studies that used wild-caught or F1 flies (e.g., Endler *et al*. 2018, Machado *et al*. 2019) or large population cages (e.g., Schou *et al*. 2017) were more likely to retain DNA viruses (figshare repository 10.6084/m9.figshare.14161250 S7).

Based on our detection thresholds, none of the public datasets we examined appeared to contain *Drosophila* Mauternbach nudivirus, *Drosophila* Yalta entomopoxvirus, the filamentous virus, the Hytrosavirus, or the three adintoviruses (figshare repository 10.6084/m9.figshare.14161250 S7). This is consistent with their extreme rarity in our own sampling, and the possibility that *Drosophila* Mauternbach nudivirus and the adintoviruses may actually infect species other than *D. melanogaster*. Although some reads from Dros-RTEC run SRR3939056 (99 flies from Athens, Georgia; Machado *et al*. 2019) did map to an adintovirus, these reads actually derive from a distinct virus that has only 82% nucleotide identity to *Drosophila*-associated adintovirus-1. Unfortunately, this closely-related adintovirus cannot corroborate the presence of *Drosophila*-associated adintovirus-1 in *D. melanogaster*, as run SRR3939056 is contaminated with *Scaptodrosophila latifasciaeformis*, which could be the host.

One of our rare viruses was present (but rare) in public data: *Drosophila* Viltain densovirus appeared only once in 3003 sequencing datasets, in one of the 63 libraries from Dros-RTEC project PRJNA308584 (Machado *et al*. 2019). *Drosophila* Tomelloso nudivirus, which was rare in our data, was more common in public data, appearing in 5 of 28 projects and 23 of 3003 runs. However, this may explained by its presence in multiple runs from each of a small number of experimental studies (e.g., Liu and Secombe 2015, Siudeja *et al*. 2015, Fang *et al*. 2017, Riddiford *et al*. 2020). Our three most common viruses were also the most common DNA viruses in public data. *Drosophila* Linvill Road densovirus appeared in 10 of the 28 projects we examined, including 363 of the 3003 runs. This virus was an exception to the general rule that DNA viruses tend to be absent from inbred or long-term laboratory lines, as it was detectable in 166 of 183 sequencing runs of inbred *D. simulans* (Signor *et al*. 2017). *Drosophila* Kallithea nudivirus appeared in four of the 28 projects, including 60 of the runs, and was detectable in wild collections of both *D. melanogaster* and *D. simulans*. *Drosophila* Vesanto virus was detectable in eight of the 28 projects, including 208 of the runs, but only in *D. melanogaster* datasets.

The presence of *Drosophila* Vesanto virus segments in public data is of particular value because it could help to elucidate patterns of segment co-occurrence. This virus was highly prevalent in a large experimental evolution study using caged populations of *D. melanogaster* derived from collections in Denmark in 2010 (Schou *et al*. 2017), where segments S01, S02, S04, S05 and S10 were almost always present, S03, S06, S07 and S08 were variable, and S09, S11 and S12 were always absent. However, because these data were derived from restriction associated digest (RAD) sequencing, absences may reflect absence of the restriction sites. *Drosophila* Vesanto virus also appeared in Pooled genome-wide association study datasets (e.g., Endler *et al*. 2018), for which segments S09 and S12 were always absent and segments S03, S10 and S11 were variable (figshare repository 10.6084/m9.figshare.14161250 S7), and in several Dros-RTEC datasets (Machado *et al*. 2019) in which only S12 was consistently absent. Unfortunately, it is difficult to test among the competing hypotheses using pooled sequencing of wild-collected flies or large cage cultures. This is because different flies in the pool may be infected with different viruses or with viruses that have a different segment composition, and because a more complex microbiome may be present. However, we were able to find one dataset from an isofemale line, GA10 collected in Athens, Georgia (USA) in 2009, that had been maintained in the laboratory for at least five generations prior to sequencing (Supporting File S6; ERR705977 from Bergman and Haddrill 2015). From this dataset we assembled 8 of the 12 segments, including two segments encoding PolB-like proteins and two encoding the DUF3472 protein. Mapping identified no reads at all from segments S9 or S12. This most strongly supports a single virus with a variable segment composition between infections and/or re-assortment. Moreover, the low species complexity of this laboratory dataset supports *D. melanogaster* as the host, with over 98% of reads mapped, and with *Drosophila*, *Wolbachia* and *Lactobacillus plantarum* the only taxa present in appreciable amounts. Example Vesanto virus sequences from these datasets are provided in figshare repository 10.6084/m9.figshare.14161250 S6.

### Genetic diversity varies among viruses and populations

We examined genetic variation in three of the most common viruses; *Drosophila* Kallithea nudivirus, *Drosophila* Linvill Road densovirus and *Drosophila* Vesanto virus. After masking regions containing indels, and using a 1% global minor allele frequency (MAF) threshold for inclusion, we identified 923 single nucleotide polymorphisms (SNPs) across the total global *Drosophila* Kallithea nudivirus pool, and 15132 distinct SNPs summed across the 44 population samples. Of these SNPs, 13291 were private to a single population, suggesting that the vast majority of *Drosophila* Kallithea nudivirus SNPs are globally and locally rare and limited to one or a few populations. This is consistent with many of the variants being recent and/or deleterious, but could also reflect a large proportion of sequencing errors—despite the analysis requiring a MAF of 1% and high base quality. Synonymous pairwise genetic diversity in the global pool was very low, at πs = 0.15%, with π at intergenic sites being almost identical (0.14%). Diversity did not vary systematically around the virus genome (figshare repository 10.6084/m9.figshare.14161250 S9). Consistent with the large number of low-frequency private SNPs, average within population-pool diversity was 10-fold lower still, at TT_S_ = 0.04%, corresponding to a very high *F*_ST_ of 0.71. In general, the level of constraint on virus genes appeared low, with global TT_A_/TT_S_ 0.39 and local TT_A_/TT_S_ = 0.58. These patterns of diversity are markedly different to those of the host, in which πS (at fourfold degenerate sites) is on the order of 1% with TT_A_/TT_S_ (zero-fold and four-fold) around 0.2, and differentiation approximately *F*_ST_ = 0.03 (Tristan *et al*. 2019, Kapun *et al*. 2020). Given that large dsDNA virus mutation rates can be 10-100 fold higher than animal mutation rates (Duffy 2018), the overall lower diversity in *Drosophila* Kallithea nudivirus is consistent with bottlenecks during infection and the smaller population size that corresponds to a 2.1% prevalence. The very low within-population diversity and high *F_ST_* and TT_A_/TT_S_ may be indicative of local epidemics, or a small number of infected hosts within each pool (expected to be 1.47 infected flies in an infected pool, assuming independence) with relatively weak constraint. Alternatively, high *F_ST_* and TT_A_/TT_S_ may indicate a high proportion of sequencing errors.

In *Drosophila* Vesanto virus we identified 4059 SNPs across all segments and divergent segment haplotypes in the global pool, with 5491 distinct SNPs summed across all infected population samples, of which 4235 were private to a single population. This corresponded to global and local diversity that was around 7-fold higher than *Drosophila* Kallithea nudivirus (global TT_S_ = 1.16%, local TT_S_ = 0.28%), and to intermediate levels of constraint on the protein sequence (TT_A_/TT_S_ = 0.20), but a similar level of differentiation (*F_ST_* = 0.76). Although the prevalence of *Drosophila* Vesanto virus appears to be slightly higher than *Drosophila* Kallithea nudivirus (2.9% vs. 2.1%), much of the difference in diversity is probably attributable to the higher mutation rates of ssDNA viruses (Duffy 2018). The apparent difference in the allele frequency distribution between these two viruses is harder to explain (73% of SNPs detectable at a global MAF of 1%, versus only 6% in *Drosophila* Kallithea nudivirus), but could be the result of the very strong constraint on protein coding sequences keeping non-synonymous variants below the 1% MAF threshold even within local populations. It is worth noting that the difference between *Drosophila* Vesanto virus and *Drosophila* Kallithea nudivirus in TT_A_/TT_S_ and the frequency of rare alleles argues against their being purely a result of sequencing error in *Drosophila* Kallithea nudivirus, as the error rates would be expected to be similar between the two viruses,

In *Drosophila* Linvill road densovirus, which was only present in 13 populations and has the smallest genome, we identified 178 SNPs across the global pool, and 253 distinct SNPs summed across the infected populations, of which 209 were private to a single population. Although this virus appears at least 6-fold less prevalent than *Drosophila* Kallithea nudivirus or *Drosophila* Vesanto virus, it displayed relatively high levels of genetic diversity both globally and locally (global TT_S_ = 1.45%, local TT_S_ = 0.21%, *F_ST_* = 0.86), and higher levels of constraint on the protein sequence (TT_A_/TT_S_ = 0.10). Given a mutation rate that is likely to be similar to that of *Drosophila* Vesanto virus, this is hard to reconcile with a prevalence that is 6-fold lower. However, one likely explanation is that *Drosophila* Linvill Road densovirus is more prevalent in the sister species *D. simulans* (above), and the diversity seen here represents rare spill-over and contamination of some samples with that species.

### Structural variation and transposable elements in Drosophila Kallithea nudivirus

*De novo* assembly of *Drosophila* Kallithea nudivirus from each sample resulted in 52 populations with complete single-scaffold genomes that ranged in length from 151.7 kbp to 155.9 kbp. Alignment showed these population-consensus assemblies to be co-linear with a few short duplications of 10-100 nt, but generally little large-scale duplication or rearrangement. Two regions were an exception to this: that spanning positions 152,180 to 152,263 in the circular reference genome (between putative proteins AQN78547 and AQN78553; genome KX130344.1), and that spanning 67,903 to 68,513 (within putative protein AQN78615). The first region comprised multiple repeats of around 100 nt and assembled with lengths ranging from 0.2 to 3.6 kbp, and the second comprised multiple repeats of around 140 nt and assembled with lengths between 0.5 and 2.4 kbp. Together, these regions explained the majority of the length variation among the Kallithea virus genome assemblies. We also sought to catalogue small-scale indel variation in Kallithea virus by analysing indels within reads. In total, after indel-realignment using GATK, across all 44 infected samples we identified 2289 indel positions in the *Drosophila* Kallithea nudivirus genome that were supported by at least 5 reads. However, only 195 of these indels were at high frequency (over 50% of samples). As would be expected, the majority (1774) were found in intergenic regions (figshare repository 10.6084/m9.figshare.14161250 S9).

Pooled assemblies can identify structural variants that differ in frequency among populations, but they are unlikely to identify rare variants within populations, such as those caused by TE insertions. TEs are commonly inserted into large DNA viruses, and these viruses have been proposed as a vector for interspecies transmission of TEs (Gilbert *et al*. 2016, Gilbert and Cordaux 2017). In total, we identified 5,169 read pairs (across 16 datasets with >300-fold coverage of *Drosophila* Kallithea nudivirus) that aligned to both *D. melanogaster* TEs and *Drosophila* Kallithea nudivirus. However, the vast majority of these (5,124 out of 5,169) aligned internally to TEs, more than 5 bp away from the start or end position of the TE, which is inconsistent with insertion (Gilbert *et al*. 2016, Loiseau *et al*. 2020). Instead, this pattern suggests PCR-mediated recombination, and assuming that all chimeras we found were artefactual, their proportion among all reads mapping to the Kallithea virus (0.01%) falls in the lower range of that found in other studies (Peccoud *et al*. 2018). We therefore believe there is no evidence supporting *bona fide* transposition of *D. melanogaster* TEs into genomes of the *Drosophila* Kallithea nudivirus in these natural virus isolates. This is in striking contrast to what was found in the *Autographa californica* multiple nucleopolyhedrovirus (Loiseau *et al*. 2020) and could perhaps reflect the tropism of *Drosophila* Kallithea nudivirus (Palmer *et al*. 2018) to tissues that experience low levels of transposition.

### A genomic insertion of Galbut virus is segregating in D. melanogaster

The only RNA virus we identified among the DNA reads from DrosEU collections was Galbut virus, a segmented and bi-parentally vertically-transmitted dsRNA *Partitivirus* that is extremely common in *D. melanogaster* and *D. simulans* (Webster *et al*. 2015, Cross *et al*. 2020). Based on a detection threshold of 0.1% of fly genome copy number, Galbut virus reads were present in 43 out of 167 samples. There are two likely sources of such DNA reads from an RNA virus in *Drosophila*. First, reads might derive from somatic circular DNA copies that are created as a part of the immune response (Mondotte *et al*. 2018, Poirier *et al*. 2018). Second, reads might derive from a germline genomic integration that is segregating in wild populations (i.e., an Endogenous Viral Element, or EVE; Katzourakis and Gifford 2010, Tassetto *et al*. 2019). We sought to distinguish between these possibilities by *de novo* assembly of the Galbut sequences from high copy-number DrosEU samples and public *D. melanogaster* DNA datasets.

We assembled the Galbut virus sequence from the three DrosEU samples in which it occurred at high read-depth: BY_Bre_15_13 (Brest, Belarus), PO_Gda_16_16 (Gdansk, Poland), and PO_Brz_16_17 (Brzezina, Poland). We were also able to assemble the sequence from four publicly available sequencing runs: three (SRR088715, SRR098913 and SRR1663569) that we believe are derived from global diversity line N14 (Grenier *et al*. 2015) collected in The Netherlands in 2002 (Bochdanovits and de Jong 2003), and SRR5762793, which was collected in Italy in 2011 (Mateo *et al*. 2018). In every case, the assembled sequence was an identical 1.68 kb, near full-length, copy of Galbut virus segment S03, including the whole of the coding sequence for the viral RNA-dependent RNA polymerase. Also, in every case, this sequence was inserted into the same location in a 297 Gypsy-like LTR retrotransposon (i.e., identical breakpoints), around 400 bp from the 5’ end. This strongly suggests that these Galbut sequences represent a unique germline insertion: Even if the insertion site used in the immune response were constant, the inserted virus sequence would be highly variable across Europe over 14 years. The sequence falls among extant Galbut virus sequences (Figure 6B), and is 5% divergent (18.5% synonymous divergence) from the closest one available in public data. The sequences are provided in figshare repository 10.6084/m9.figshare.14161250 S10

**Figure 6:**
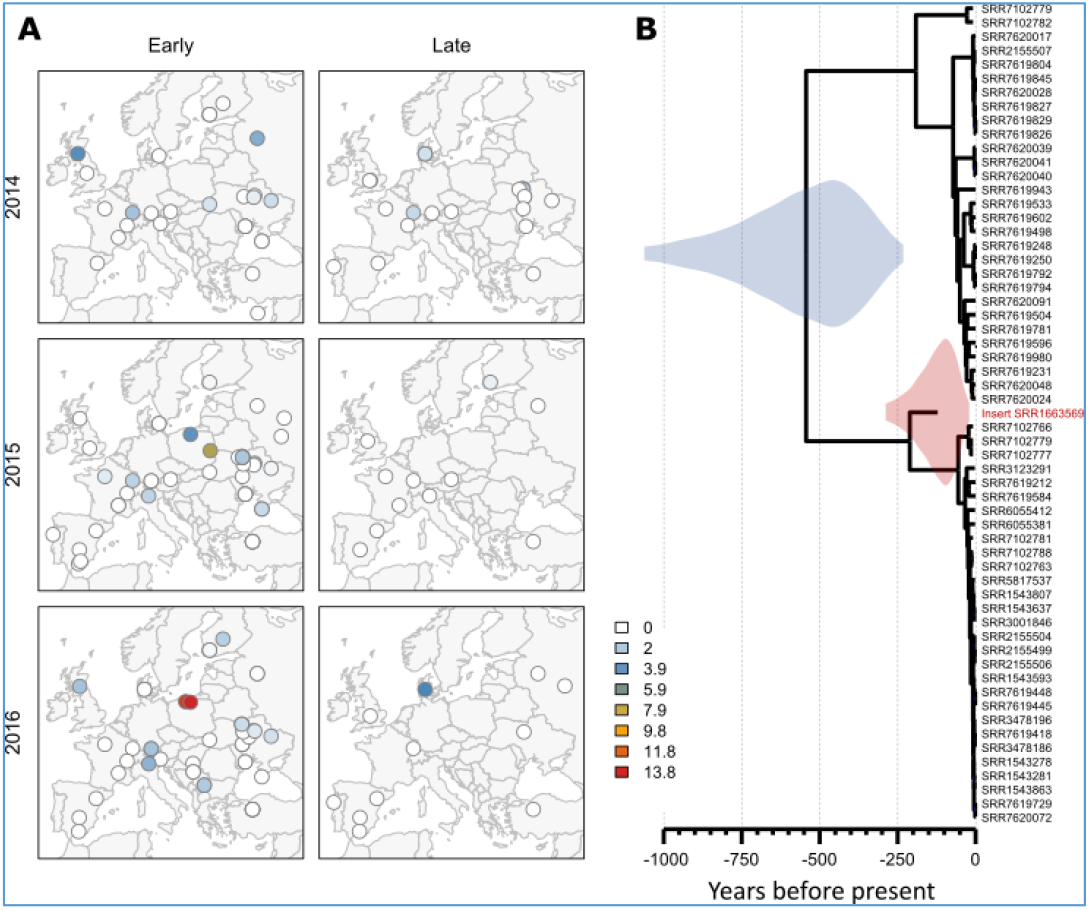
*Drosophila melanogaster* harbours an endogenous genomic copy of Galbut virus. (A) Maps show the spatial distribution of the DNA reads from the Galbut EVE, as a percentage of fly genomes (maximum 13.8%) on colour scale. Rows show years of sampling, and columns show ‘early’ or ‘late’ samples in each year (B) The relationship between the Galbut EVE and Galbut virus sequences detectable in public datasets, illustrated by a Bayesian maximum clade-credibility tree inferred under a strict clock, with median-scaled node dates. The 95% highest posterior density for the root date of extant Galbut viruses is shown in blue (230-1060 years before present), and the 95% highest posterior density for the inferred date of insertion, is shown in red (20-290 years before present)

Interestingly, populations with a substantial number of Galbut virus reads (a maximum of 13.8% or 11 chromosomes of 80) appeared geographically limited, appearing more commonly in higher latitudes, and with a different spatial distribution in the early and late collecting seasons (ΔDIC = 26.92; Figures 5C, 6A). Given the absence of this sequence from Dros-RTEC (Machado *et al*. 2019), DGRP (Mackay *et al*. 2012) and the other *Drosophila* Genome Nexus datasets (Lack *et al*. 2015, Lange *et al*. 2016), it seems likely that this insertion is of a recent, likely northern or central European, origin. We used a strict-clock phylogenetic analysis of viral sequences to estimate that the insertion occurred within the last 300 years (posterior mean 138 years ago, 95% highest posterior density interval 20-287 years ago; Figure 6B), i.e. after *D. melanogaster* was spreading within Europe. Unfortunately, the insertion site in a high copynumber transposable element means that we were unable to locate it in the genome. This also means that it was not possible to detect whether the insertion falls within a piRNA-generating locus, which is seen for several endogenous viral elements (EVE) in mosquitoes (Palatini *et al*. 2017) and could perhaps provide resistance to the vertically transmitted virus. Surprisingly, DNA reads from Galbut virus were more likely to be detected at sites with a higher percentage of reads mapping to *Wolbachia* (95% credible interval for the effect [0.074,0.41]; ΔDIC = −5.52). Given that no correlation between Galbut virus and *Wolbachia* has been detected in the wild (Webster *et al*. 2015, Shi, White, *et al*. 2018), we think this most likely reflects a chance association between the geographic origin of the insertion and the spatial distribution of *Wolbachia* loads (Kapun *et al*. 2020).

## Discussion

Although metagenomic studies are routinely used to identify viruses and virus-like sequences (e.g., Shi, Zhang, *et al*. 2018, Zhang *et al*. 2018), simple bulk sequencing can only show the presence of viral sequences; it cannot show that the virus is replicating or transmissible, nor can it unequivocally identify the host (reviewed in Obbard 2018). This behoves metagenomic studies to carefully consider any additional evidence that might add to, or detract from, the claim that an ‘associated virus-like sequence’ is indeed a virus. A couple of the DNA viruses described here undoubtedly infect *Drosophila. Drosophila* Kallithea nudivirus has been isolated and studied experimentally (Palmer *et al*. 2018), and *Drosophila* Tomelloso nudivirus is detectable in some long-term laboratory cultures (e.g. Liu and Secombe 2015, Siudeja *et al*. 2015, Fang *et al*. 2017, Riddiford *et al*. 2020). Others, such as *Drosophila* Viltain densovirus, *Drosophila* Linvill Road densovirus, and *Drosophila* Vesanto virus, are present at such high copy numbers, and sometimes in laboratory cultures, that any host other than *Drosophila* seems very unlikely. Some, appearing at reasonable copy number but in a single sample, could be infections of contaminating *Drosophila* species (*Drosophila* Mauternbach nudivirus, the adintoviruses), or spill-over from infections of parasitoid wasps *(Drosophila* Yalta entomopoxvirus, the filamentous virus). A few, having appeared at low copy number in a single sample, could be contaminants—although we excluded virus-like sequences that appeared strongly associated with contaminating taxa (figshare repository 10.6084/m9.figshare.14161250 S4).

These caveats aside, along with *Drosophila innubila* nudivirus (Unckless 2011) and Invertebrate iridescent virus 31 in *D. obscura* and *D. immigrans* (Webster *et al*. 2016), our study increases the total number of published DNA viruses associated with *Drosophila* to sixteen. Although a small sample, these viruses hint at some interesting natural history. First, it is striking that more than a third of the reported DNA viruses are Nudiviruses (six of the 16 published, plus a seventh from *Phortica variegata*; Figure 2). This suggests that members of the Nudiviridae are common pathogens of *Drosophila*, and may indicate long-term host lineage fidelity with short-term switching among species. Such switching is consistent with the lack of congruence between host and virus phylogenies, and the fact that both *D. innubila* Nudivirus and *Drosophila* Kallithea Nudivirus infect multiple *Drosophila* species (Figure 2). Second, the majority of DNA viruses seem to be rare. Seven of the twelve viruses confidently ascribable to *D. melanogaster* or *D. simulans* were detected in just one of the 167 population samples, and likely only one of 6668 flies, consistent with a European prevalence less than 0.07%. Only *Drosophila* Vesanto virus and *Drosophila* Kallithea nudivirus seem relatively common—being detected in more than half of populations and having estimated prevalences of 2.9% and 2.1%, respectively. It is unclear why DNA viruses should have such a low prevalence, on average, as compared to RNA viruses (Webster *et al*. 2015). In simple ‘susceptible-infected-susceptible’ compartment models, low pathogen prevalence can result from high lethality, low transmission rates, or high recovery rates (relative to baseline mortality rates). It is therefore possible that DNA virus infections are less persistent than RNA virus infections or that they have lower transmission rates. Alternatively, this may reflect sampling bias, such that DNA viruses increase morbidity to the extent that infected flies are less likely to be sampled than uninfected flies. This may also explain why DNA viruses rarely persist through multiple generations in laboratory fly lines. Alternatively, it may be that the rare viruses represent deadend spill-over from other taxa that can only be seen here because of the large sample size. Third, although some viruses showed broad geographic patterns in prevalence, a lack of repeatability associated with sampling location, and the very high *F_ST_* values, hint that transient local epidemics may be the norm, with viruses frequently appearing and then disappearing from local fly populations.

Finally, *Drosophila* do indeed seem to harbour fewer DNA viruses than RNA viruses, supporting an observation that was made before any had been described (Brun and Plus 1980, Huszart and Imler 2008). This cannot simply be an artefact of reduced sampling effort, as almost all *Drosophila*-associated viruses have been reported from undirected metagenomic studies, and metagenomic studies of RNA are as capable of detecting expression from DNA viruses as they are of detecting RNA viruses (e.g., Webster *et al*. 2015). Instead, it suggests that the imbalance must reflect some aspect of host or virus biology. For example, it may be a consequence of differences in prevalence. If RNA viruses have higher prevalence in general, or specifically in those adult flies attracted to baits, and/or RNA viruses persist more easily in fly or cell cultures, then this may explain their more frequent detection.

Taken together, our analyses of the distribution and diversity of DNA viruses associated with *Drosophila melanogaster* at the pan-European scale provide an ecological and evolutionary context for studies of host-virus interaction in *Drosophila*. However, we currently lack almost any data on the natural host range or fidelity of *Drosophila* viruses, and we have no knowledge of their real-world fitness consequences for the host. In the future, such information will be vital if we are to capitalise on *Drosophila* models to understand the co-evolutionary process.

## Acknowledgements

We thank all of the members of the DrosEU and Dros-RTEC communities for their ongoing engagement in this collaborative European project. We are especially grateful to the teachers Antonio J. Buendía, Ma Josefa Gómez, Ma Luisa Espinosa and the students of the IES Eladio Cabañero (Tomelloso, Spain), and to the teachers David González, Silvana Castillo and the students of the IES José de Mora (Baza, Spain), who contributed to fly collections in 2016 as part of the “Melanogaster Catch the Fly” citizen science project. We thank Alex Twyford for providing computing time, and Lewis Stevens and Andrew Rambaut for their help with MinION sequencing.

## Funding

Megan Wallace was supported by the UK Natural Environmental Research Council through the E3 doctoral training programme (NE/L002558/1), and Sanjana Ravindran was supported by Wellcome Trust PhD programme (108905/Z/15/Z).

Andrea Betancourt received funding from BBSRC grant BB/P00685X/1

Thomas Flatt received funding from Swiss National Science Foundation grants 31003A-182262, PP00P3_165836, and PP00P3_133641/1.

Clément Gilbert received funding from Agence Nationale de la Recherche (grant ANR-15-CE32-0011-01)

Josefa González received funding from the European Research Council (ERC) under the European Union’s Horizon 2020 research and innovation programme (H2020-ERC-2014-CoG-647900) and from the Fundación Española para la Ciencia y la Tecnologia-Ministerio de Economía y Competitividad (FCT-15-10187).

Sonja Grath received funding from Deutsche Forschungsgemeinschaft grant GR 4495/2

Maaria Kankare received funding from Academy of Finland projects 268214 and 322980.

Martin Kapun received funding from Austrian Science Fund (FWF) grant P32275.

Volker Loeschcke received funding from Danish Research council for natural Sciences (FNU) grant nr 4002-00113B

Banu Sebnem Onder received funding from the Scientific and Technological Research Council of Turkey (TUBITAK) (Grant No. 214Z238)

John Parsch received funding from Deutsche Forschungsgemeinschaft grant PA 903/8

Marina Stamenkovic-Radak, Marija Savic Veselinovic, and Mihailo Jelic received funding from the Ministry of Education, Science and Technological Development of the Republic of Serbia (Grant number 451-03-68/2020-14/200178)

Fabian Staubach received funding from Deutsche Forschungsgemeinschaft grant STA1154/4-1; Projektnummer 408908608

Marija Tanaskovic, Aleksandra Patenkovic, and Katarina Eric received funding from the Ministry of Education, Science and Technological Development of the Republic of Serbia (Grant number 451-03-68/2020-14/200007)

The DrosEU consortium has been funded by a Special Topics Network (STN) grant by the European Society of Evolutionary Biology (ESEB).

Files available via figshare 10.6084/m9.figshare.14161250

S1: Excel spreadsheet detailing collection dates and locations

S2: Text document listing the microorganisms including in the ‘Drosophila microbiome’ mapping reference, and the mitochondrial and plastid sequences included in the species diagnostic mapping reference.

S3: Excel spreadsheet detailing the mapped read numbers from the DrosEU data. Sheet A gives raw mapped read counts, Sheet B gives counts normalised to read length in reads per kilobase per million reads (RPKM), Sheet C gives raw counts of reads mapping to additional Species-diagnostic loci.

S4: DNA Fasta file of virus fragments thought to be associated with contaminating taxa

S5: Excel spreadsheet detailing the presence and read counts of DNA viruses in DrosEU datasets. Sheet A gives counts normalised to the fly to give virus copy number in genomes fly genome and estimated prevalence at three different detection thresholds, Sheet B provides metadata used for the statistical analysis.

S6: DNA fasta file of assembled *Drosophila* Vesanto virus segments, including divergent segments and segments assembled from public datasets.

S7: Excel spreadsheet detailing the presence and read counts of DNA viruses in 28 publicly available *Drosophila* sequencing projects. Sheet 1 summarises the public datasets included, Sheet 2 gives raw mapped read counts

S8: Excel spreadsheet detailing mean and total TT_A_, TT_S_ and TT_A_/TT_S_ for each gene (sheet A) and the number of synonymous and non-synonymous SNPs in the genome of *Drosohila* Kallithea nudivirus, *Drosophila* Linvill Road densovirus and *Drosophila* Vesanto virus (sheet B).

S9: Figure showing A) variation in nucleotide diversity across non-coding and synonymous sites in the *Drosophila* Kallithea nudivirus genome, plotted as a sliding window with two window sizes, and B) the percentage of *Drosophila* Kallithea nudivirus infected samples that showed evidence of an indel. Intergenic regions of the genome are coloured in grey. A chi-square test for independence found a strong positive association between intergenic regions and InDels (X-squared = 3236, df = 1, p-value < 2.2e^-16^).

S10: DNA fasta file of exemplar Galbut virus sequences aligned with the EVE and Gypsylike LTR retroelement 297.

## Notes

https://doi.org/10.6084/m9.figshare.14161250.v1

### Competing Interest Statement

The authors have declared no competing interest.

