## Supplementary figures and images for "The discovery, distribution and diversity of DNA viruses associated with *Drosophila melanogaster* in Europe" (from another version of this preprint: version 1, 2020-10-16)

### Figure1.pdf

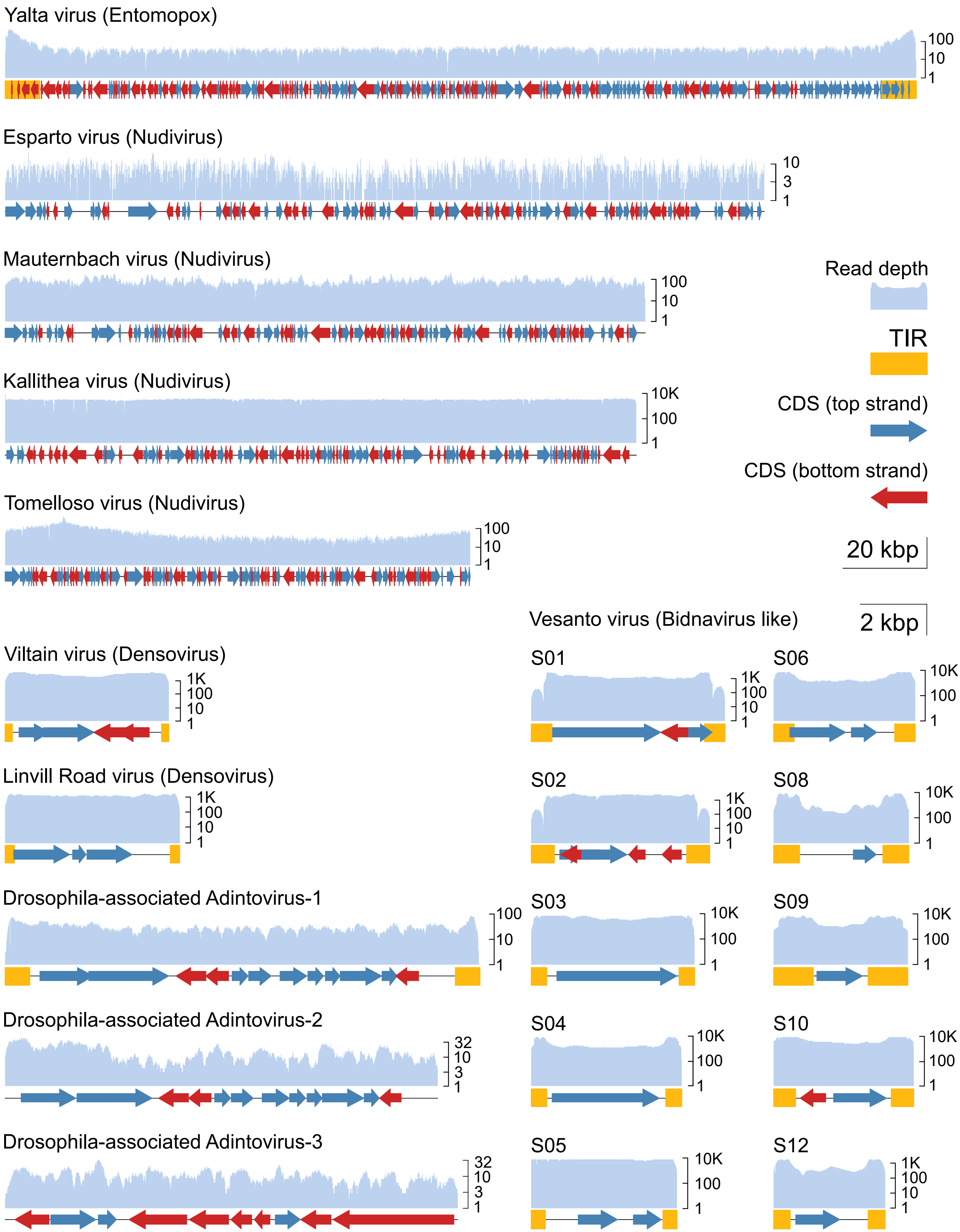

### Figure2.pdf

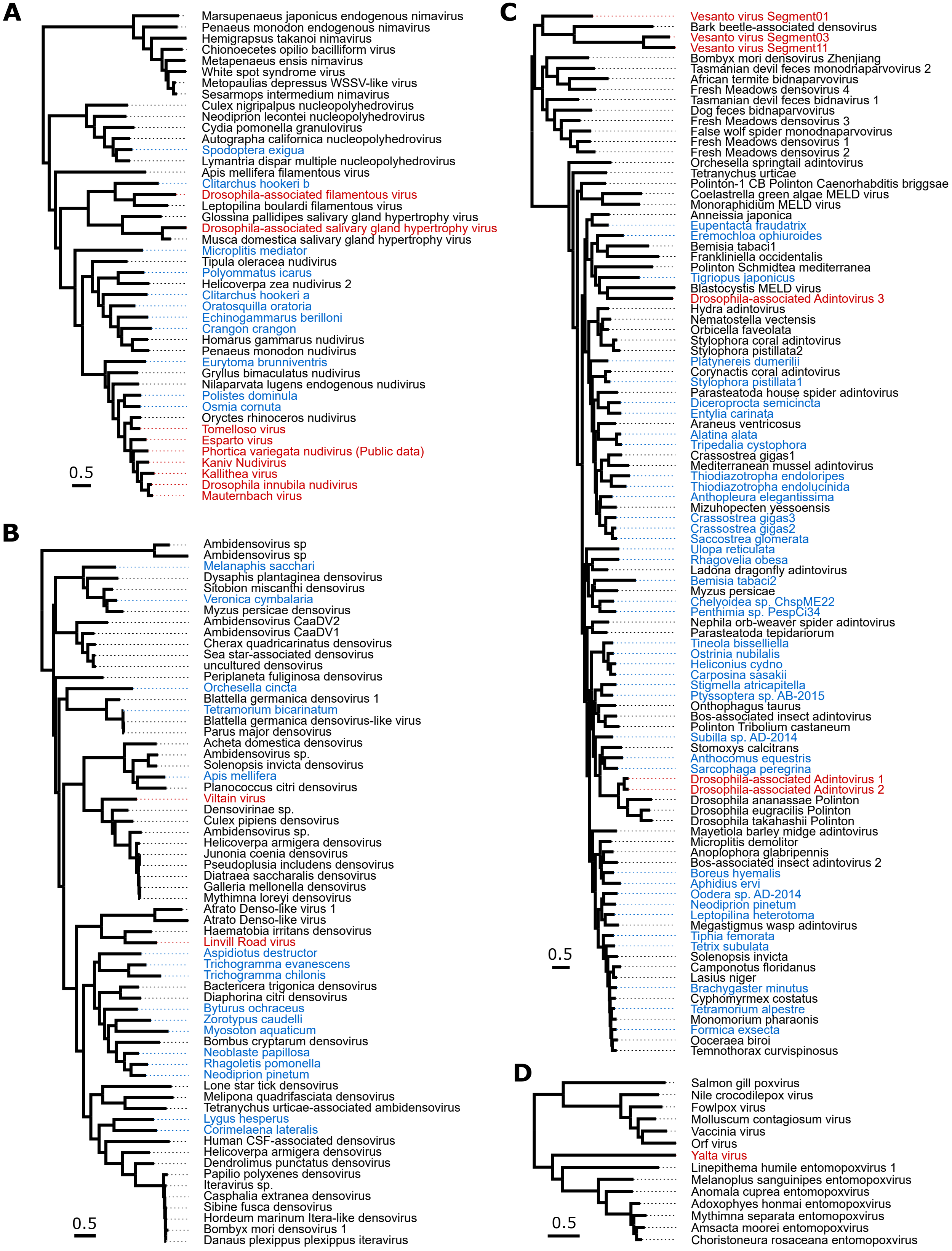

### Figure3.pdf

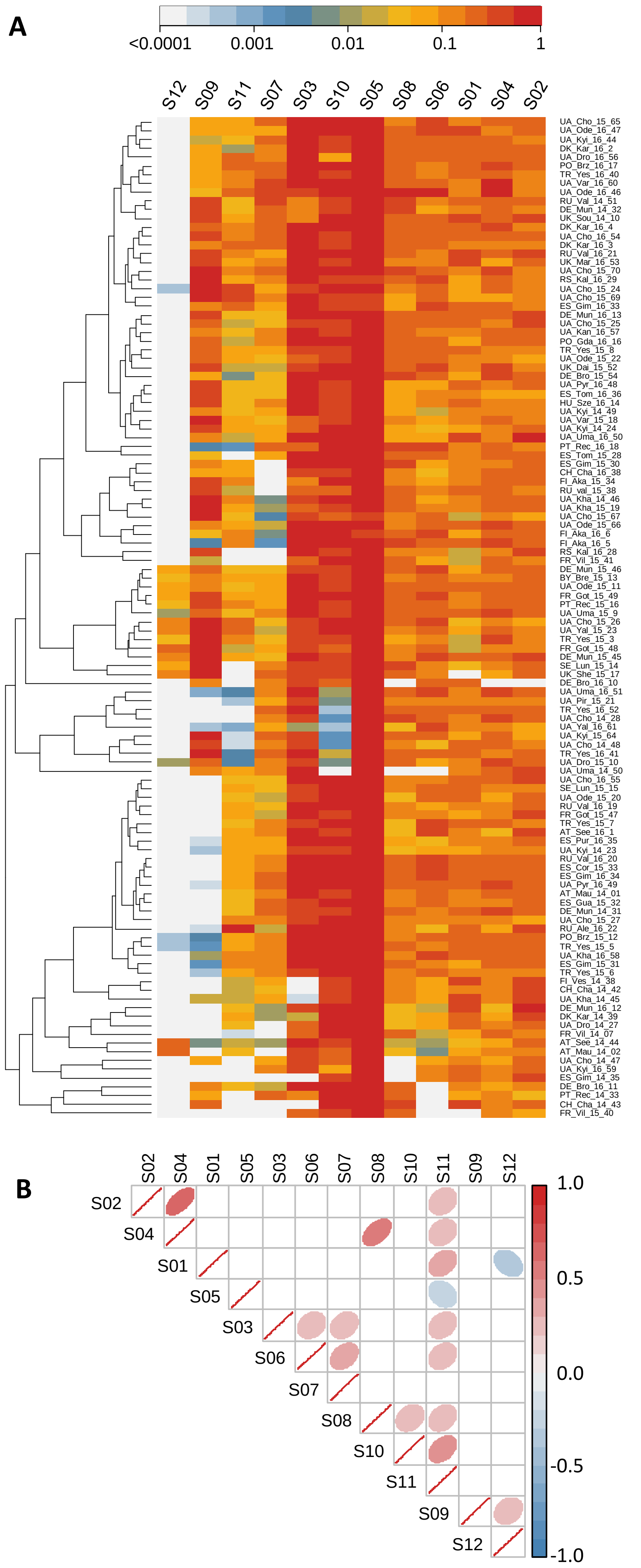

### Figure4.png

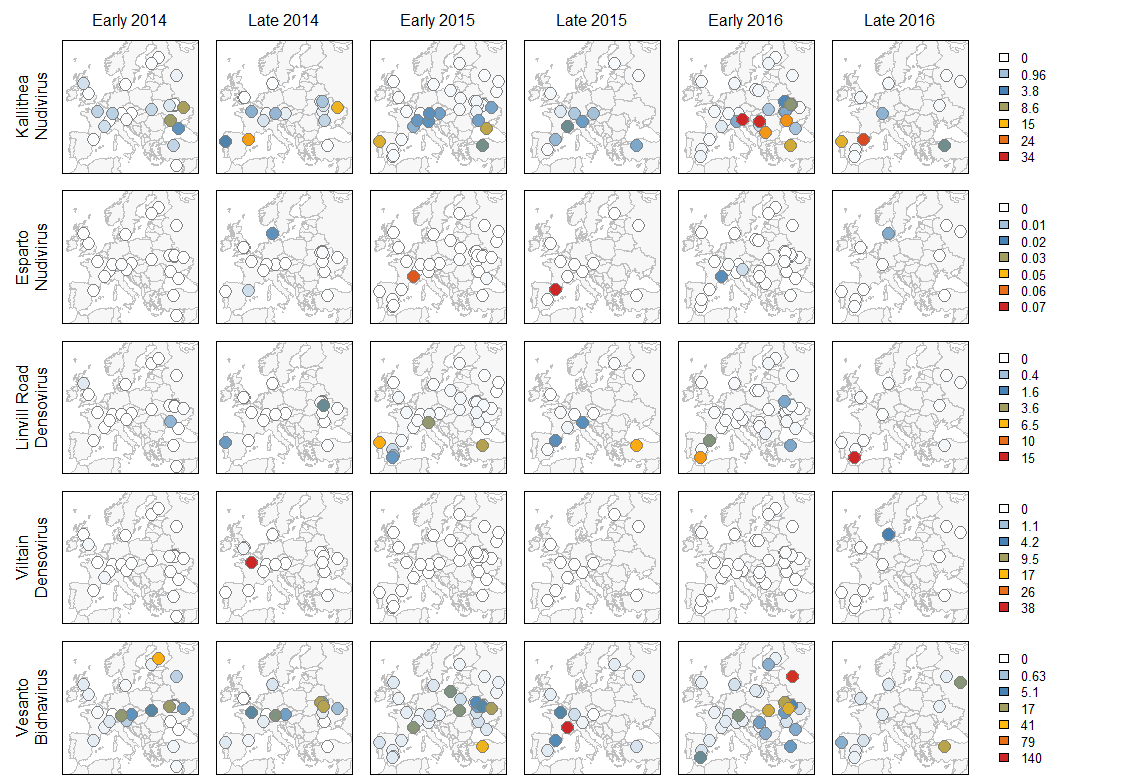

### Figure5.pdf

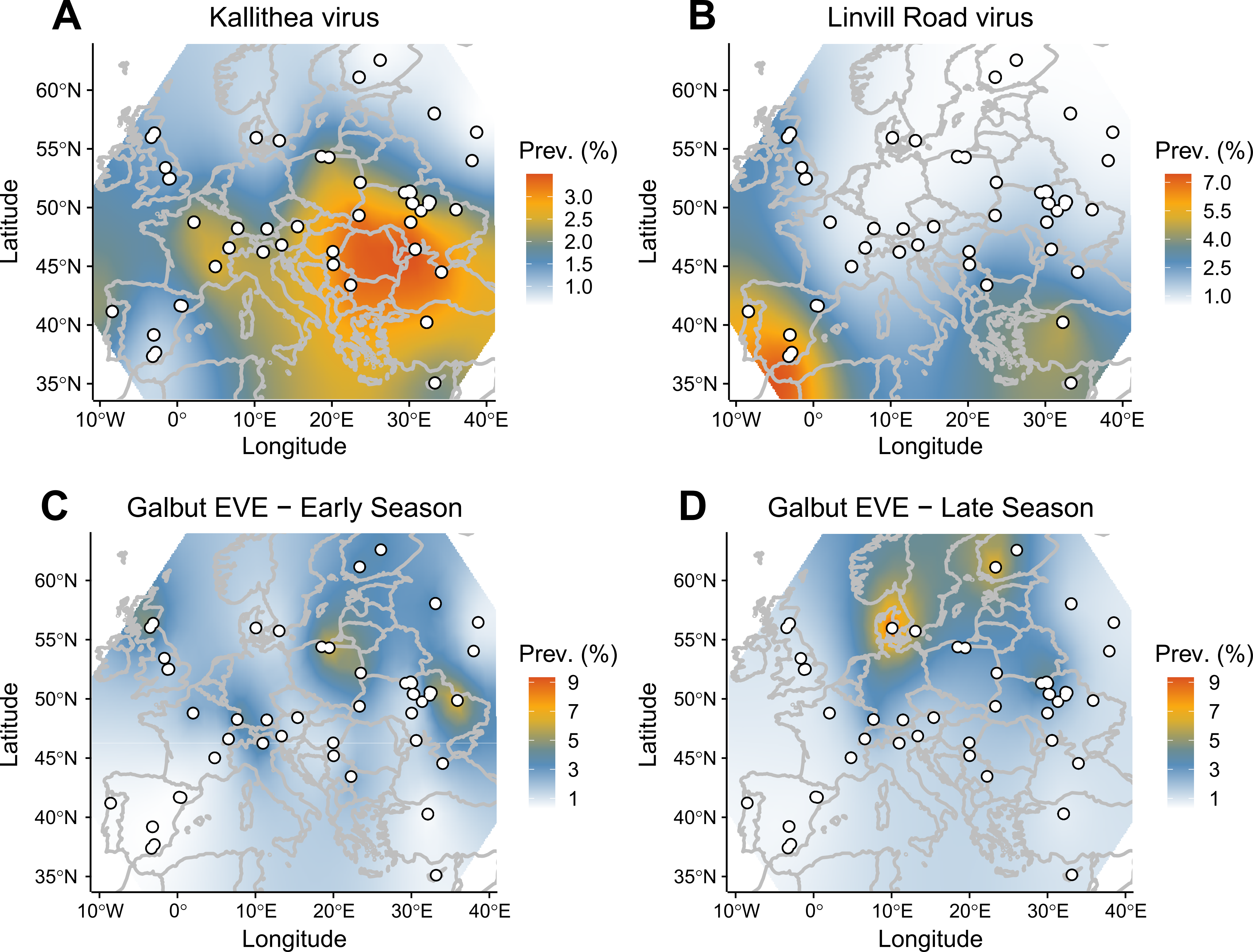

### Figure6.pdf

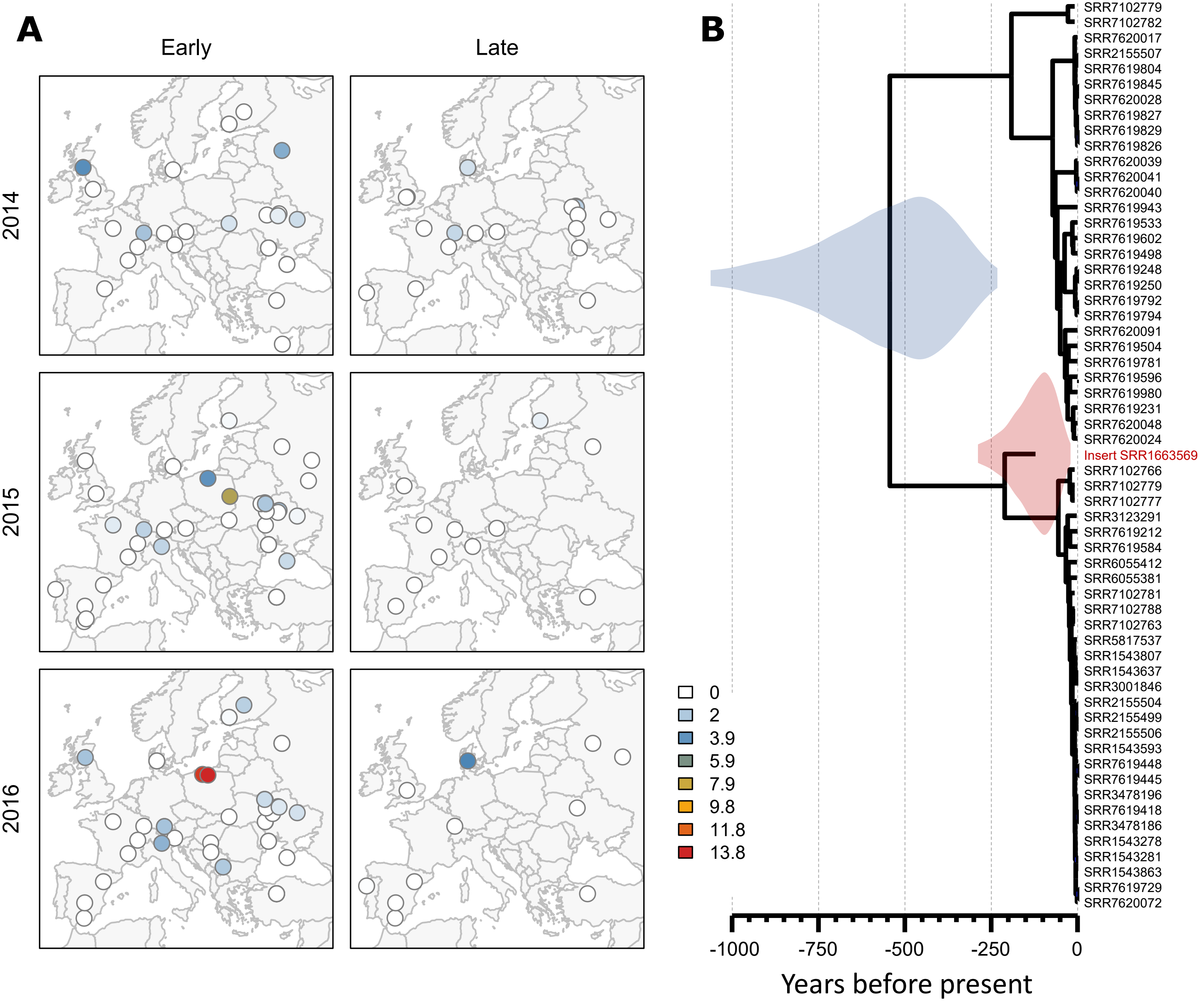

### Supplementary_File_S9.pdf

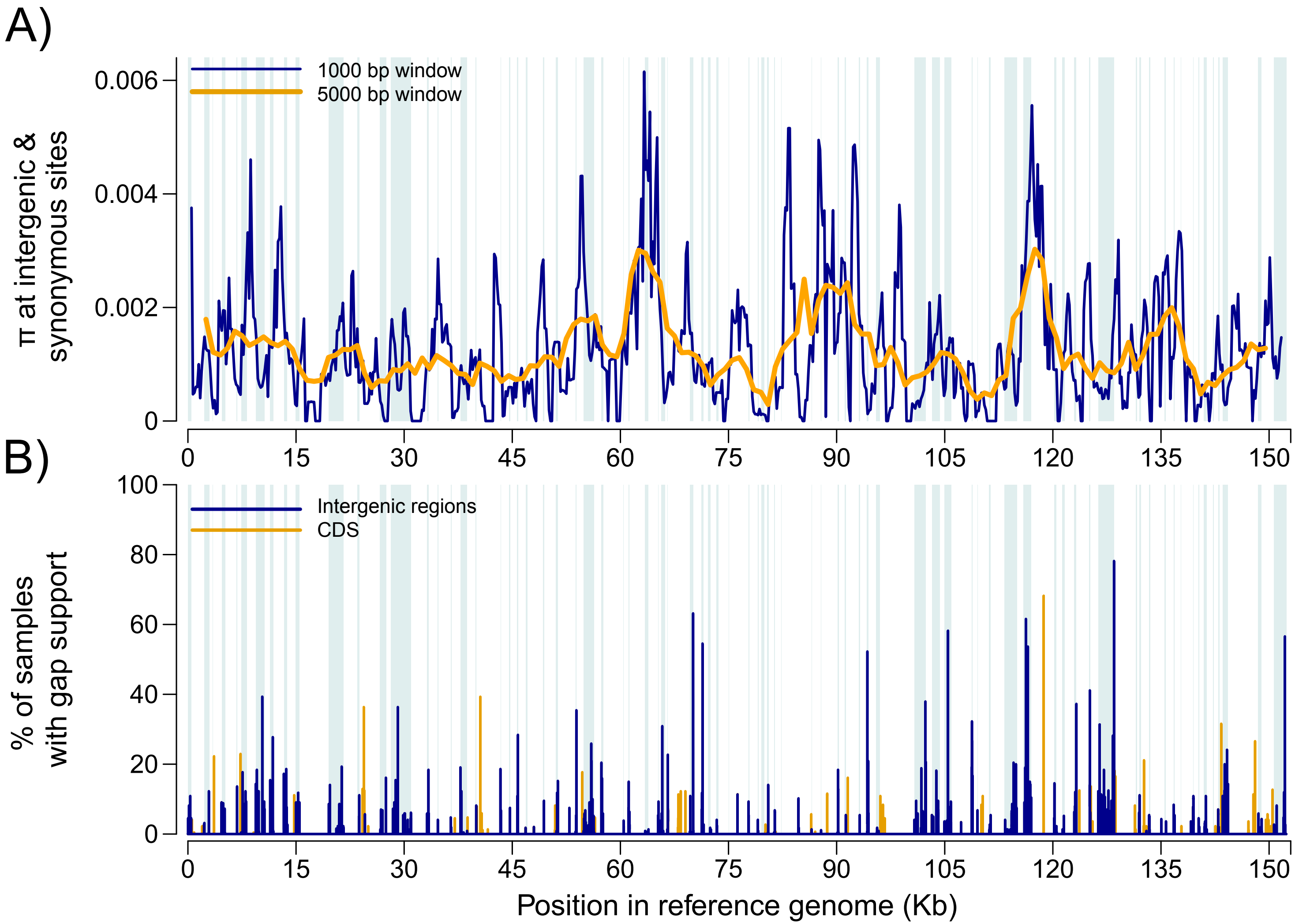
